# A meta-analysis of chromatin-associated loci provides insights into mechanistic interpretations of trait heritability

**DOI:** 10.64898/2026.03.19.712994

**Authors:** Max F. Dudek, Brandon M. Wenz, Benjamin F. Voight, Laura Almasy, Struan F.A. Grant

**Affiliations:** Center for Spatial and Functional Genomics, Children’s Hospital of Philadelphia, Philadelphia, PA 19104, USA; Graduate Group in Genomics and Computational Biology, Perelman School of Medicine, University of Pennsylvania, Philadelphia, PA 19104, USA; Genetics and Epigenetics Program, Cell and Molecular Biology Graduate Group, Biomedical Graduate Studies, University of Pennsylvania - Perelman School of Medicine, Philadelphia, PA, USA; Department of Genetics, Perelman School of Medicine, University of Pennsylvania, Philadelphia, PA 19104, USA; Department of Systems Pharmacology and Translational Therapeutics, University of Pennsylvania, Perelman School of Medicine, Philadelphia, PA 19104, USA; Institute for Translational Medicine and Therapeutics, Perelman School of Medicine, University of Pennsylvania, Philadelphia, PA 19104, USA; Corporal Michael J. Crescenz VA Medical Center, Philadelphia, PA, USA; Lifespan Brain Institute, Children’s Hospital of Philadelphia and Perelman School of Medicine, University of Pennsylvania, Philadelphia, PA, USA; Department of Biomedical and Health Informatics, Children’s Hospital of Philadelphia; Department of Pediatrics, Perelman School of Medicine, University of Pennsylvania, Philadelphia, PA 19104, USA; Division of Human Genetics, Children’s Hospital of Philadelphia, Philadelphia, PA 19104, USA; Division of Endocrinology and Diabetes, Children’s Hospital of Philadelphia, Philadelphia, PA 19104, USA

## Abstract

The vast majority of trait-associated loci discovered through genome-wide association studies (GWAS) are non-coding, yet most lack statistical alignment with any discovered expression quantitative trait loci (eQTLs). In particular, eQTLs are depleted at gene-distal regions and at “functionally important” genes – those with strong selective constraint and complex regulatory landscapes – likely due to selective depletion of high-effect variants. Here, we investigate the role of variants with weaker effects on expression transmitted through distal regulatory elements, which are detectable as chromatin accessibility QTLs (caQTLs). We aggregated caQTL data from ten studies derived across different tissues, cell-types and lines, representing 104,024 lead caQTLs across 3,457 samples. We found that, across a range of gene properties, caQTLs are discovered at functionally important genes more often than eQTLs. These observations are consistent with a model in which many eQTLs and GWAS hits are mediated through genetic effects on regulatory elements, which may have weak or context-dependent effects on gene expression. Our results suggest that caQTL discovery is more sensitive than eQTL discovery in capturing the molecular consequences of GWAS hits, and can provide complimentary information to eQTLs by implicating functional mechanisms of additional disease-associated loci.

## Introduction

Almost 90% of trait-associated variants revealed by GWAS are harbored within non-coding genomic regions^1,2^ making it challenging to elucidate the specific genetic mechanisms by which each variant contributes to complex physiologic traits and disease. Many GWAS signals are linked to gene regulatory elements and mechanistically modulate physiological traits through variation that also changes gene expression, (i.e., eQTLs), suggesting that they act principally through *cis*-regulatory mechanisms^3–6^. However, at current sample sizes, most GWAS hits are not immediately attached to an eQTL-like signal that might provide a mechanistic explanation^7^; indeed, an analysis of human gene expression across 49 tissue types by the Genotype-Tissue Expression (GTEx) Consortium estimated that only 43% of GWAS hits statistically linked to (i.e., colocalized with) an eQTL signal^6^. Many factors have been hypothesized to contribute to this “colocalization gap”, including insufficient statistical power^5,8,9^, a lack of eQTL discovery in the correct context^7^ (e.g. a specific developmental stage^10^, cell type^11^, or cell state in response to a stimulus^12^), and post-transcriptional effects (e.g. splicing) not mediated through expression levels.

In a previous report, Mostafavi et al.^13^ noted that at current sample sizes, GWAS signals (i.e. GWAS “hits”) are biased towards different classes of genes from eQTLs. Specifically, compared to eQTLs, GWAS hits are (1) more distal to genes, and (2) more likely to be found near genes with high selective constraint, complex regulatory landscapes, and high connectedness in co-expression networks. The depletion of eQTLs at these “functionally important” genes suggests that in addition to uncovering eQTLs in more contexts and larger sample sizes, there is scope for additional types of molecular assays to further close the colocalization gap.

In particular, the compaction or openness of chromatin is a central determinant of gene transcription, both at promoters and at more distal *cis*-regulatory elements (cREs), whose activity can enhance or repress the expression of nearby genes^14–16^. One popular approach to identify cREs leverages the Assay for Transposase-Accessible Chromatin using sequencing (ATAC-seq)^17^, which defines peaks of open chromatin and their relative accessibility. Genetic variants associated with change in chromatin peak accessibility are known as chromatin accessibility QTLs (i.e., caQTLs)^18^. Consistent with the belief that the genetic association of eQTLs is often mediated through their effect on cREs, genetic variants explain a greater proportion of heritability with chromatin accessibility than with gene expression, and are more readily discovered as caQTLs than eQTLs^19–23^. In addition, caQTLs colocalize with more GWAS hits than eQTLs when discovered in the same set of samples^23–25^. caQTLs can also capture genetic effects on enhancer priming across a broad range of dynamic cell states, while their downstream effects on gene expression are only detectable as eQTLs if profiled in the correct cellular context^26^. However, the relationship between caQTL and eQTL discoverability and their mechanistic linkage to GWAS hits at different types of genes is not currently well understood.

To explore in more detail, we assembled a large collection of caQTLs currently available in the public domain to examine their potential mechanistic role in physiologic trait association compared directly against eQTLs. We hypothesize that the limited power in eQTL discovery is due to weaker or context-dependent perturbations on expression mediated through cREs, which caQTL discovery would be more sensitive in capturing. Initially, we replicate the previous comparison and observation between eQTLs and GWAS hits from Mostafavi et al., expanding this effort to include an intermediate step: genetic variation changes chromatin accessibility which, in turn, modulates cis-gene expression, then ultimately to the downstream physiologic trait of study. We collected caQTLs from ten studies^21,22,24,27–33^ and found that, compared to discovered eQTLs, the properties of caQTLs more closely resemble those of GWAS hits. Specifically, caQTLs are less depleted than eQTLs near “functionally important” genes – those with strong selective constraint and complex regulatory landscapes. We show that these observations are consistent with a model in which many eQTLs and GWAS hits are mediated through genetic effects on cREs. This model implies that for such GWAS hits, caQTLs are better powered than eQTLs to detect colocalizations due to a more direct genetic effect. In particular, the model expects caQTLs to colocalize with GWAS hits more in distal regions and at “functionally important” genes compared to eQTLs, helping to explain what we observe. These results suggest that with a limited sample size for discovery, epigenetic association signals can provide complimentary information to eQTLs by implicating functional mechanisms of additional disease-associated loci.

## Results

### Comparison of caQTLs to eQTLs and GWAS hits

Following replication of the analysis of eQTLs and GWAS hits carried out by Mostafavi et al., we incorporated a collection of single nucleotide polymorphism (SNP) caQTL summary statistics gathered from ten different studies across multiple cell and tissue types (**Fig. 1A**, **Supplementary Table 1**). We ran linkage-disequilibrium (LD) clumping to identify independent lead caQTLs, then removed any SNPs in strong LD (r^2^>0.8) with protein-altering SNPs. In order to compare caQTLs to GTEx eQTLs, we also removed SNPs further than 1 Mb from a transcription start site (TSS), in line with what was not tested by GTEx for eQTLs; this resulted in a list of 104,024 caQTLs. We then ran this catalog of caQTLs through the computational pipeline utilized by Mostafavi et al., allowing us to compare eQTLs, caQTLs, and GWAS hits directly (**Methods**). Briefly, we computed average values for a range of SNP and gene properties in each list of caQTLs, eQTLs, and GWAS hits. For gene properties, we first assigned every SNP to its nearest gene (creating lists of “GWAS genes”, “eQTL genes”, and “caQTL genes”, **Fig. 1B**), and considered the properties of that gene. To account for potential confounding effects due to differences between caQTLs, eQTLs, and GWAS hits, we also ran this analysis on lists of control SNPs for caQTLs, eQTLs, and GWAS hits, matched for minor allele frequency (MAF), LD score, and gene density (**Fig 1C**). See **Methods** for full details.

**Figure 1.**
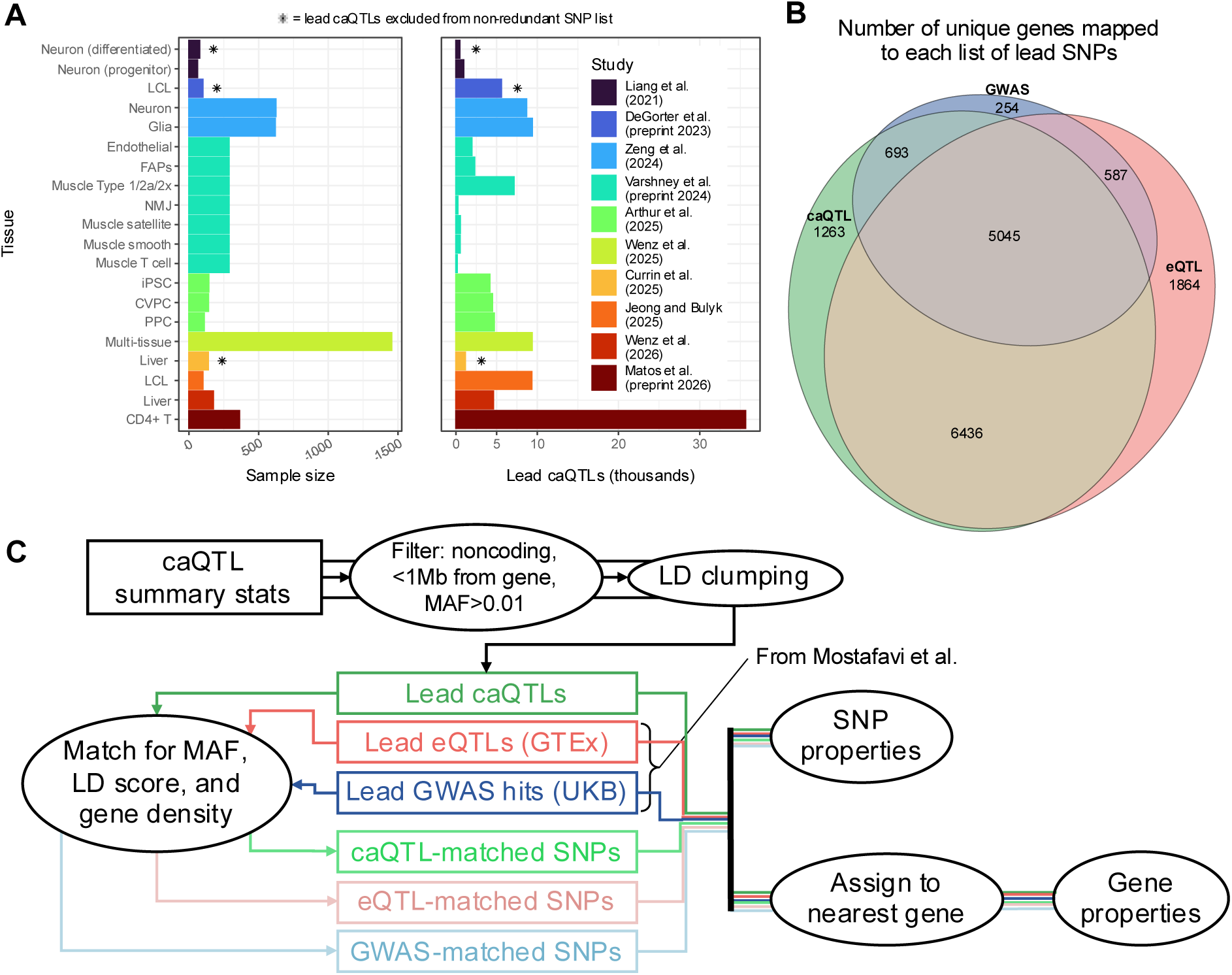
A meta-analysis of caQTL discovery. (A) Details of studies used in this analysis. The number of lead caQTLs reported are those remaining after LD clumping and filtering. Redundant tissues that were excluded from the aggregate analyses are marked with a star. Full details and statistics for studies used can be found in **Supplementary Table 1**. (B) Venn diagram showing which genes mapped (by nearest TSS) to at least one of each type of SNP. (C) Schematic of SNP analysis. caQTL summary stats from multiple studies were filtered and clumped separately, and the resulting lists of lead caQTLs were concatenated into one list.

### caQTLs are less depleted around functionally important genes than eQTLs

SNPs with large effect sizes on complex traits have been shown to have lower minor allele frequencies due to natural selection^34–37^. The degree to which a gene is constrained by selection (a proxy for “functional importance”) can be measured by the pLI (probability of being loss-of-function intolerant) score^38^, with previous studies having shown that SNP-disease heritability is enriched near high-pLI genes^38–40^. However, Mostafavi et al. and other studies^38,41–43^ reported that such highly constrained genes are *depleted* for eQTLs, especially those with large effects, suggesting that natural selection purges SNPs which strongly affect the expression of these genes. In our replicated analysis, while GWAS hits were more frequently found near high-pLI genes (pLI>0.9) compared to control SNPs (26% vs 21%, *P*=1.2×10^−7^), the opposite was true for eQTLs (12% vs 18%, *P*=3.0×10^−34^) (**Fig. 2A**). Interestingly however, the depletion for caQTLs was far lower than for eQTLs, and not significant (20% vs 21%, *P*=0.077).

**Figure 2.**
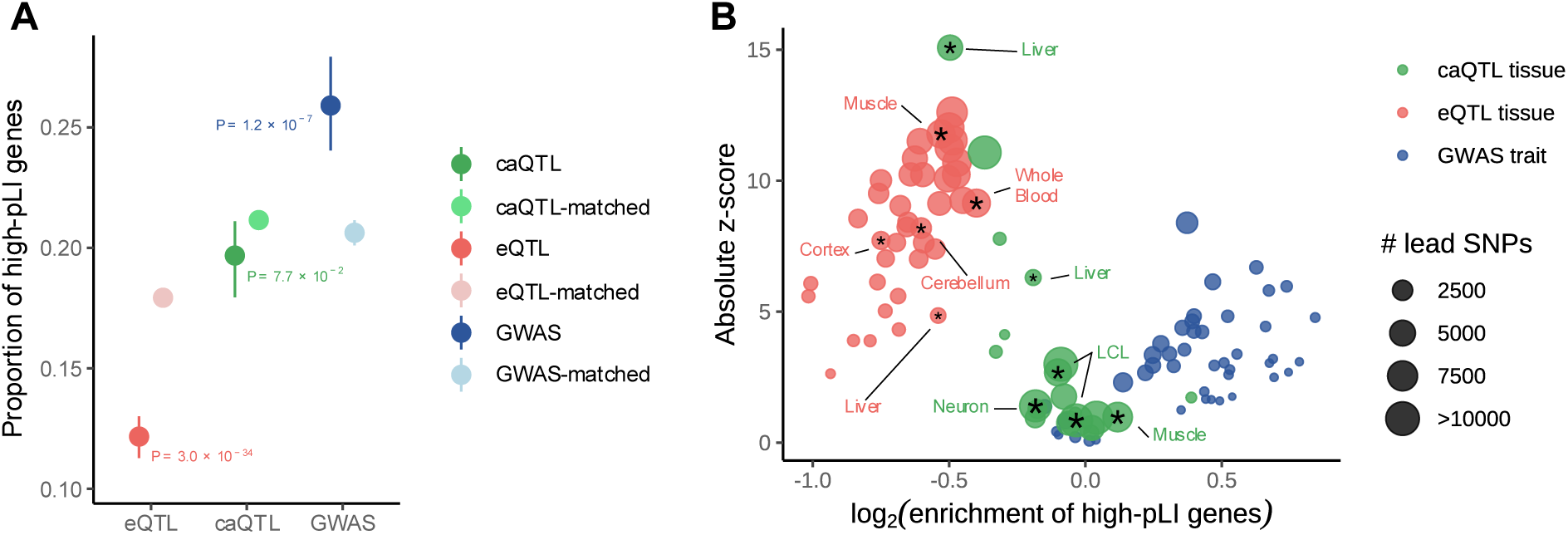
caQTLs are found more often than eQTLs at highly constrained genes. (A) Proportion of SNP-mapped genes with pLI>0.9, for each list of SNPs. Confidence intervals for matched lists show 2.5-97.5 percentiles, confidence intervals for lists of discovered SNPs are from bootstrapping. See **Methods** for *P*-value calculations. (B) Volcano plot of SNP enrichment in high-pLI (>0.9) genes, separated by tissue or trait. caQTL tissues are shown multiple times if they are represented in separate studies. For comparison, selected QTL tissues are labelled and marked with a star for clarity.

In addition to the selective constraint of genes, Mostafavi et al. also considered the level of regulatory complexity of genes near trait and expression-associated SNPs. Their analyses showed that GWAS hits were found more frequently near genes with more complex regulatory landscapes than eQTLs. Specifically, compared to genes near eQTLs, genes near GWAS hits have more TSSs, have longer enhancers, are more connected in co-expression networks, and are more likely to be transcription factors (TFs). When including caQTLs in these comparisons, we found that the regulatory complexities of caQTL genes were between those of eQTL genes and GWAS genes (**Fig. 3**). For example, when considering the number of different TSSs used by genes^44^, GWAS genes had more TSSs than genes mapped to control SNPs (6.4 vs 5.6, *P*=3.8×10^−7^), eQTL genes had fewer (4.4 vs 5.0, *P*=6.2×10^−12^), while caQTL genes did not show a significant difference (5.8 vs 5.8, *P*=0.57) (**Fig. 3A**). Mostafavi et al. calculated the total length of enhancers for every gene, based on enhancer–gene links inferred from the Roadmap Epigenomics Consortium dataset^45^. Using logistic regressions to distinguish QTLs/GWAS genes from genes linked to random SNPs (**Methods**), we found that GWAS genes had longer enhancers (*P*=4.0×10^−5^), eQTL genes had shorter enhancers (*P*=2.5×10^−9^), while for caQTL genes they were slightly longer (*P*=1.1×10^−3^) (**Fig. 3B**). A longer set of enhancers is associated with weaker enhancer activity, since each individual peak contributes proportionally less to the expression of the gene. Thus, this result suggests that caQTLs are found in weaker enhancers compared to eQTLs.

**Figure 3.**
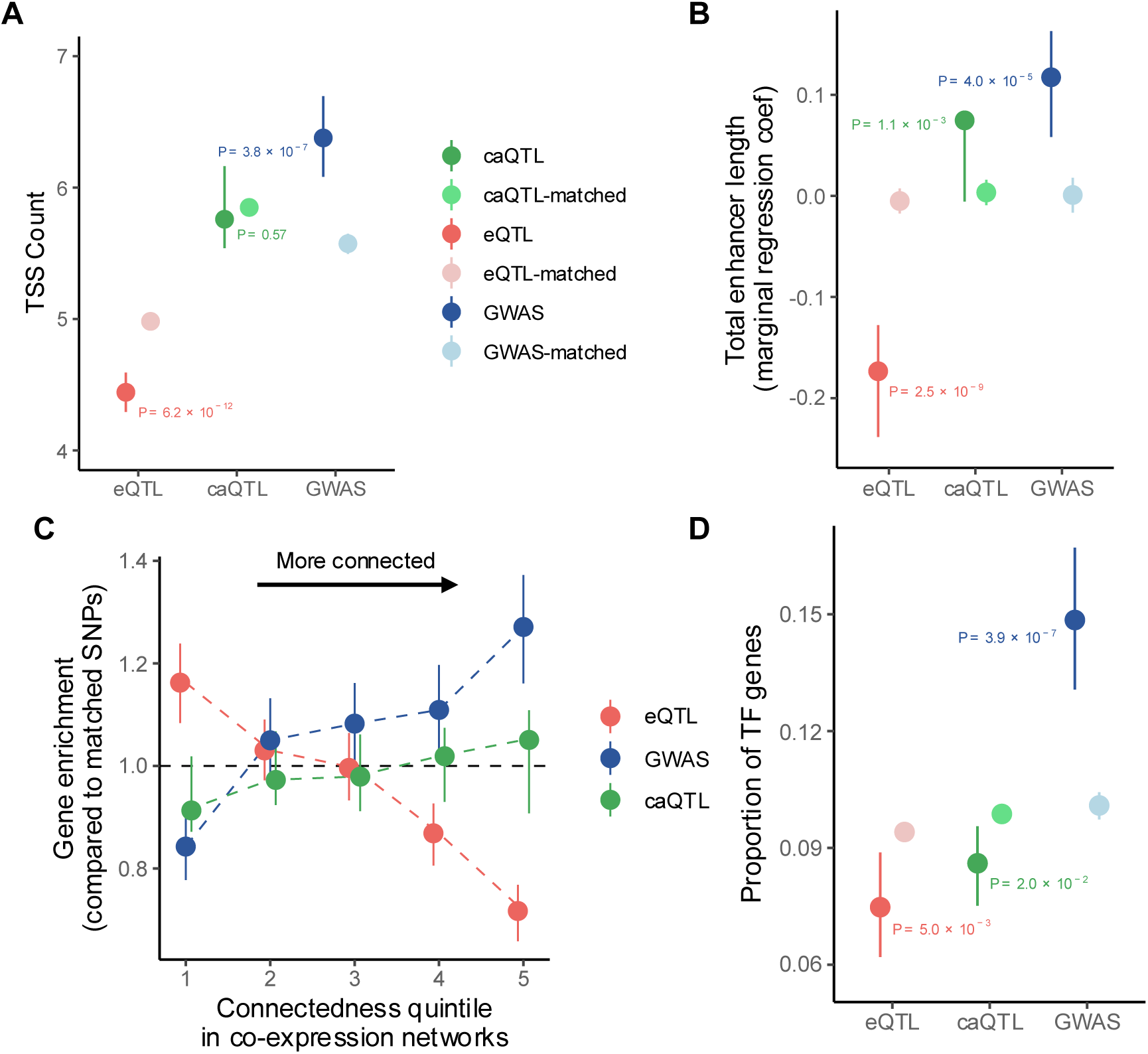
Nearest-gene properties of QTLs and GWAS hits. (A) Number of TSSs in genes mapped from QTLs/GWAS hits. (B) Marginal coefficients for “total enhancer length” in logistic regressions which distinguish QTLs/GWAS hits from random SNPs (**Methods**). Enhancer length measured using Roadmap. (C) Enrichment of genes mapped to QTLs/GWAS hits compared to genes mapped to matched SNPs, in bins of genes ranked by connectedness in co-expression networks from Saha et al.^46^ (D) Proportion of genes mapped from QTLs/GWAS hits that are transcription factors (TFs). For all panels: confidence intervals for matched lists show 2.5-97.5 percentiles, confidence intervals for lists of discovered SNPs are from bootstrapping. For the full list of calculated gene properties, see **Supplementary Table 2** and **Supplementary Figs. 1-3**. See **Methods** for *P*-value calculations.

We also considered the measure of gene connectivity in Mostafavi et al., calculated from the number of neighbors a gene had in coexpression networks inferred by Saha et al. for GTEx tissues^46^. While highly connected genes were enriched near GWAS hits and depleted near eQTLs, there was no significant enrichment of genes in the top connectivity-quintile near caQTLs compared to control SNPs (*P*=0.41) (**Fig. 3C**). Interestingly, not all properties of caQTL genes were intermediate between eQTL and GWAS genes. For example, the proportion of genes that were transcription factors (TFs) was higher for GWAS genes compared to control-SNP genes (15% vs 10%, *P*=3.9×10^−7^), while it was lower for caQTL genes (8.6% vs 9.9%, *P*=0.02), similar to eQTLs (**Fig. 3D**).

To confirm that these observed differences between caQTLs and eQTLs were not due to differences in study design, sample size, or sample cohorts, we also compared caQTLs and eQTLs which were both discovered by the Parker lab in the same set of muscle samples^29^ (**Methods**). The differences in gene properties between eQTLs and caQTLs described above remained consistent if we limited the analysis to only QTLs discovered in this set of muscle cells (**Supplementary Fig. 4**), indicating that they were not driven by differences between GTEx and caQTL discovery cohorts. Together, these results suggest that while genes with high regulatory complexity are depleted of eQTLs, they are not depleted for caQTLs.

### caQTLs are discovered at more TSS-distal loci compared to eQTLs and GWAS hits

While eQTLs are known to be strongly enriched close to TSSs^47,48^, Mostafavi et al. observed that GWAS hits are less enriched, and on average lie further away from genes than eQTLs. caQTLs were even less enriched in promoters and more distal than GWAS hits, with only 19% of caQTLs lying within 10kb of a gene compared to 13% for control SNPs (**Fig. 4A**). Additionally, caQTLs were more enriched vs matched SNPs in enhancers compared to eQTLs, particularly in distal enhancers defined by the ENCODE project^49^ (2.1-fold for caQTLs vs 1.2-fold for eQTLs, **Fig. 4B**). Given that GWAS hits also showed stronger enhancer enrichment compared to eQTLs, this suggests that current caQTLs have a higher probability of tagging an underlying molecular mechanism for disease-associated SNPs in enhancers compared to current catalogs of eQTL variation.

**Figure 4.**
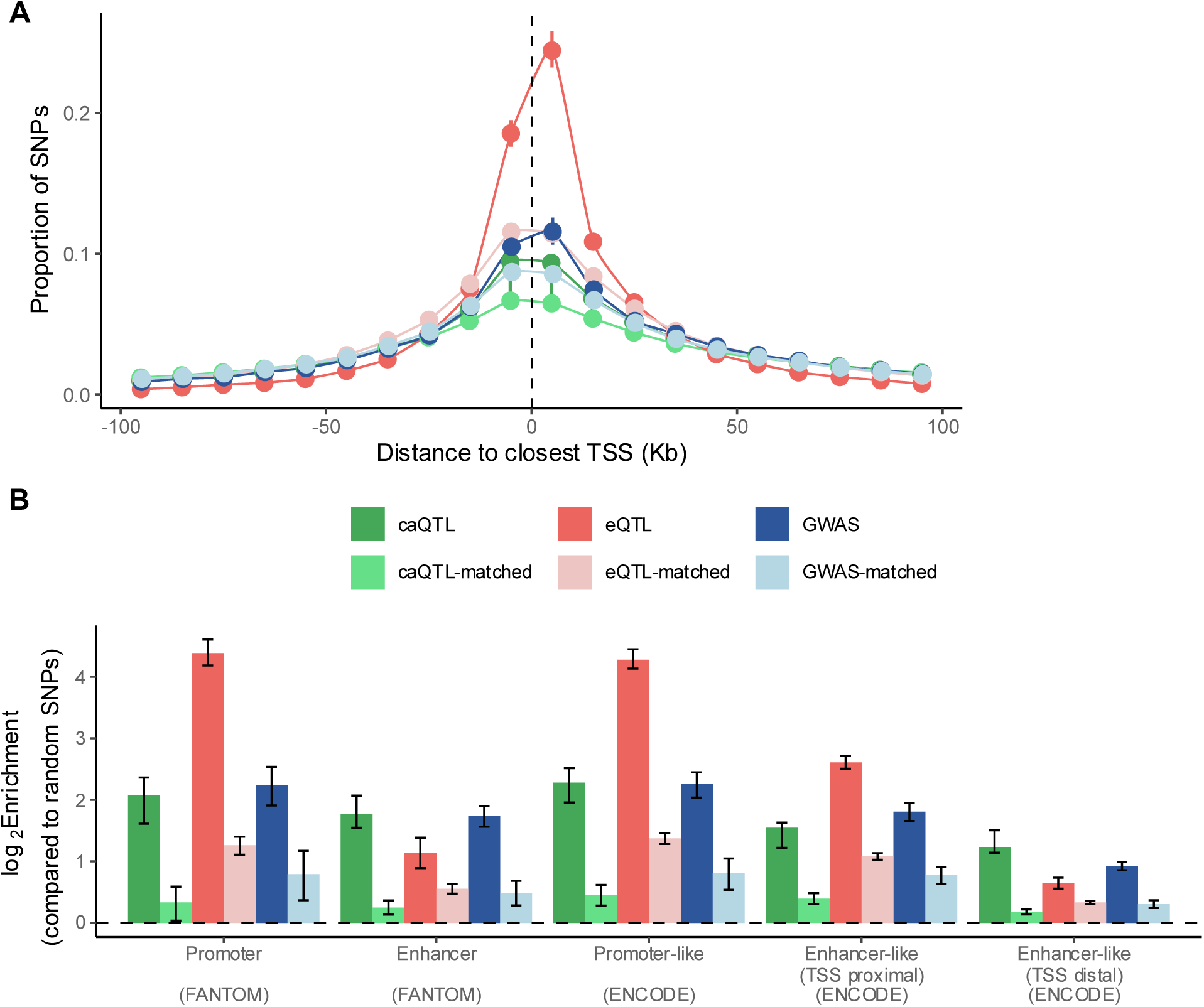
SNP properties of QTLs and GWAS hits. (A) Proportion of QTLs/GWAS hits in bins based on distance to the nearest TSS. Each bin is 10 Kb wide. SNPs >100 Kb from a TSS are not shown. (B) Enrichment of QTLs/GWAS hits in promoter and enhancer elements annotated in the FANTOM project (left), and in the ENCODE project (right). For each annotation, the enrichment value is computed as the fraction of SNPs in the annotation divided by the fraction of all SNPs in the annotation, which is then log_2_ transformed. For all panels: confidence intervals for matched lists show 2.5-97.5 percentiles, confidence intervals for lists of discovered SNPs are from bootstrapping.

### An extended model of functional variant discovery

To address the observed properties of caQTLs compared to those of eQTLs and GWAS hits, we expanded the model of discovery presented in Mostafavi et al. to include a chromatin accessibility component. Given that many genes are widely considered to be regulated through distal cREs, we elected to consider the subset of GWAS hits in which a variant directly perturbs a cRE, which in turn directly modulates the expression of a trait-relevant gene (**Methods**, **Supplementary Note**). For consistency with the Mostafavi model, we labeled the three relevant effect sizes as *β*_1,_ *β*_2,_ and *γ*. (**Fig. 5A**). We then modeled the discoverability of a single SNP as a function of these (squared) effect sizes, and visualized the range of effect sizes for which that SNP was expected to be discovered as a caQTL, eQTL, or GWAS hit. A simple model which does not account for natural selection demonstrated that a more direct genetic effect on chromatin accessibility compared to gene expression led to a wider range of SNP discovery for caQTLs compared to eQTLs, even when the caQTL sample size was lower (**Fig. 5B**). Under natural selection, this discrepancy became greater for SNPs which affect the expression of “high-trait effect” genes, or those with high *γ*^2^ values (**Figs. 5C-D**). This is consistent with our observation that caQTLs appear more frequently than eQTLs at genes that are enriched for GWAS hits.

**Figure 5.**
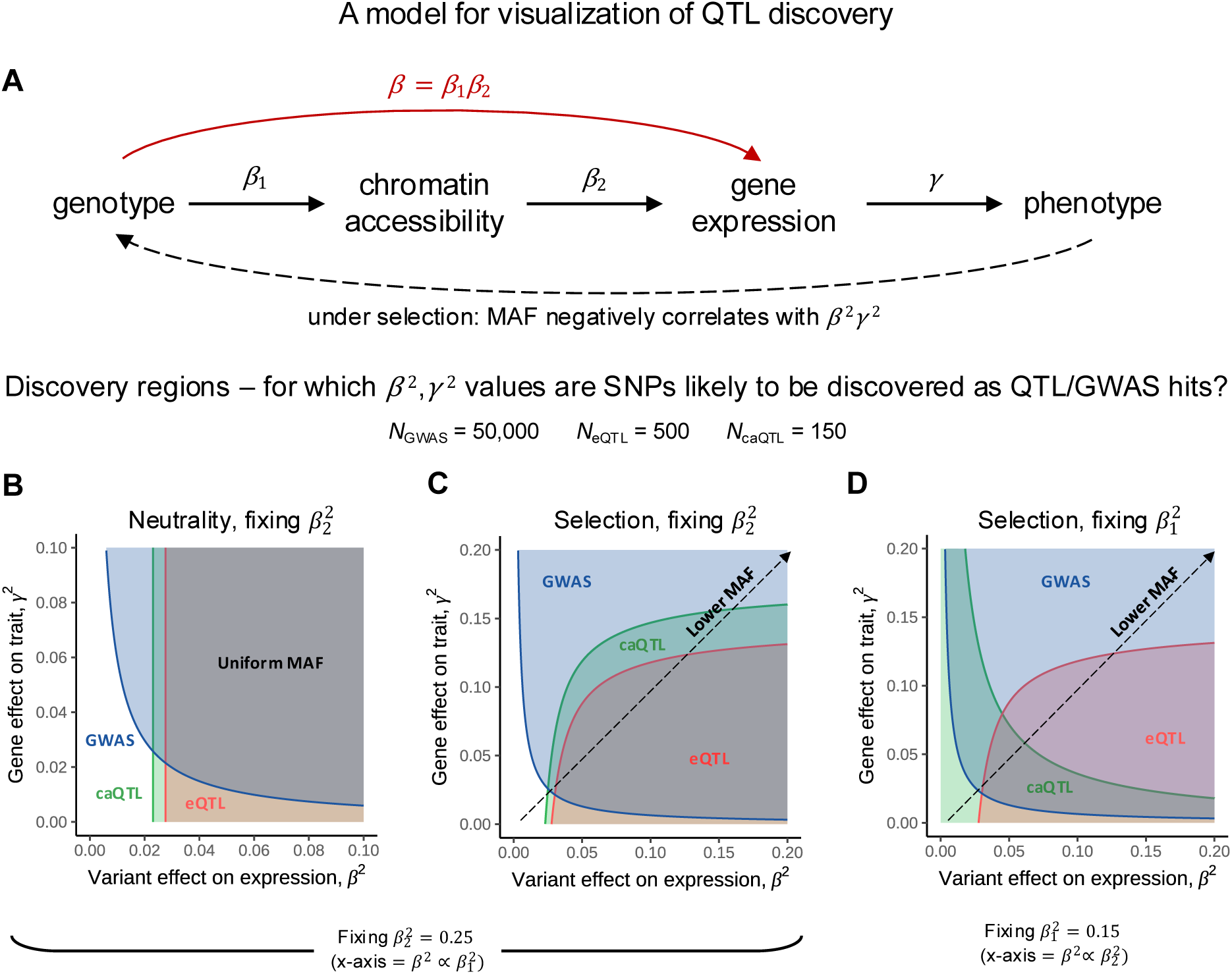
A model of QTL and GWAS discovery. (A) A simple causal model considering the effect of a single SNP on the accessibility of a single peak (or cRE), the expression of a single gene, and a single phenotypic trait. This model applies to the subset of SNPs whose effect on gene expression is mediated through chromatin accessibility. If natural selection is not modeled, the minor allele frequency (MAF) is uniform; under selection, it is negatively correlated with the effect of the SNP on the trait. (B-D) Discovery regions showing the values of *β*^2^ and *γ*^2^ for which a SNP is likely to be discovered as a caQTL (green), eQTL (red), or GWAS hit (blue). Because the model includes three effect sizes (*β*_1,_ *β*_2,_ *γ*) that are free to vary, each visualization fixes either *β*_1_^2^ or *β*_2_^2^. All plots use a GWAS sample size of 50,000, an eQTL sample size of 500, and a caQTL sample size of 150. For discovery regions under selection, κ = 750 is used to control the strength of selection. All variances are standardized so that the squared effect sizes (*β*_1_^2^, *β*_2_^2^, *γ*^2^) represent r^2^ values, the proportion of variance in the outcome explained by variance in the causal variable. (B) Discovery regions under no natural selection, where *β*_2_^2^ is fixed to be 0.25, so that *β*^2^ on the x-axis is proportional to *β*_1_^2^. (C) Discovery regions under selection, where *β*_2_^2^ is fixed to be 0.25, so that *β*^2^ on the x-axis is proportional to *β*_1_^2^. (D) Discovery regions under selection, where *β*_1_^2^ is fixed to be 0.15, so that *β*^2^ on the x-axis is proportional to *β*_2_^2^. See the **Supplementary Note** for a full explanation of the model and discovery regions, along with links to interactive plots where all parameters of the model can be adjusted.

In particular, the model showed that for smaller values of *β*_2_^2^ (the effect of accessibility on expression), caQTLs were especially more likely to be discovered alongside GWAS hits compared to eQTLs (**Fig. 5D**). The reason for this was that weaker enhancers (i.e., peaks with lower effects on expression) were less constrained by selection, and thus more likely to harbor common variants affecting accessibility. However, these variants in weaker enhancers had a lower net effect on expression (*β*^2^ = *β*_1_^2^*β*_2_^2^) and were therefore harder to discover as eQTLs. They will be discovered only as caQTLs, and if the gene has a sufficient effect on the trait (*γ*^2^), as a GWAS hit. This observation aligns strongly with a population-scale multiomic QTL discovery from FinnGen^23^. In this study, SNPs at constrained genes (in the model, those with high *γ*^2^) showed similar effects on chromatin accessibility (*β*_1_^2^) to unconstrained genes, while effects on gene expression were smaller at constrained genes and mediated through systematically weaker enhancer-gene links (smaller *β*_2_^2^). This observation also explains the relative frequency of caQTLs, eQTLs, and GWAS hits near these constrained genes.

## Discussion

Previous analyses have shown that “functionally important” genes – those with high levels of selective constraint and regulatory complexity – are enriched for GWAS hits, yet depleted of eQTLs, suggesting that natural selection reduces the allele frequency of SNPs with a high effect on expression near such genes. This hinders eQTL discovery near these genes at current sample sizes. caQTLs potentially represent SNPs with weaker effects on expression that are mediated through cREs, which should be less depleted by selection. Here, we show that caQTLs appear more frequently at functionally important genes compared to eQTLs, though not as frequently as GWAS hits. These results suggest that caQTLs have utility in interpreting GWAS hits at functionally important genes which currently have no colocalizing eQTL. Note that while caQTLs had a similar distribution of gene properties to control SNPs, they are enriched for GWAS loci and in known regulatory regions^24,28,31–33^, distinguishing them from randomly chosen background SNPs.

Our observations are consistent with a model in which the effect of a SNP on gene expression is mediated through chromatin accessibility of a cRE. Clearly, this model does not apply to all non-coding SNPs – SNPs in promoters, for instance, can modulate expression without affecting a distal enhancer at all. Furthermore, this simplified model also ignores pleiotropic effects, by which a SNP influences both expression and accessibility independently, or instances where gene transcription causes an opening of chromatin in the region. However, our results comparing caQTLs to eQTLs suggest that the accessibility to expression causal pathway dominates over alternative pathways, and indeed the outsized contribution of caQTLs to disease heritability compared to eQTLs indicates that caQTLs are the primary mediator of genetic effects on gene expression^23^. While the genetic architecture of trait association can potentially be more accurately visualized as a dense web of causal relationships between genes, regulatory elements, and other epigenetic factors (e.g. methylation, 3D interactions), such modeling is beyond the scope of this study. Instead, we propose that chromatin accessibility lies on the earlier side of this regulatory cascade and thus is more directly influenced by genetics. As a result, caQTL discovery is more sensitive to the effect of SNPs compared to eQTL discovery. We also note that this principle could generalize to other types of QTLs, such as methylation QTLs^50^ or looping QTLs^51^.

A prominent limitation of this study is the relative non-uniformity of caQTL discovery across tissues, compared to the discovery of eQTLs across tissues in GTEx, or the discovery of GWAS hits across traits in the UKBB. While discovery cohorts in GTEx or the UKBB largely overlap between tissues or traits, the caQTL studies analyzed here used separate cohorts. In addition, each caQTL discovery may employ slightly different peak calling algorithms, window sizes for tests, or models and methods used to assess significance. Furthermore, the set of tissues and cell types analyzed in caQTL studies is far less diverse and comprehensive than the set of GTEx tissues. Some tissues are represented in multiple studies, whereas many are not represented in any. However, the results that we show here appear to be largely consistent across tissue types and studies, even when caQTLs and eQTLs are discovered in the same set of samples, suggesting that those properties of caQTLs are not sensitive to tissue type or discovery parameters.

It is also important to note that the differences we describe occur for SNPs *discovered at current sample sizes*. Indeed, recent studies observed that as eQTL sample size increases, the depletion of eQTLs at constrained genes decreases^9^, and eQTLs at constrained genes are more likely to colocalize with GWAS hits^23^. Thus the observed bias is not intrinsic to different types of variants but the result of a limited sample size, and increasing eQTL power should remain an important aim in closing the colocalization gap. Instead, our results suggest that while sample size for molecular QTL studies is limited, epigenetic signals such as caQTLs can capture weaker genetic regulatory effects with less dependence on context compared to eQTLs, and can provide complimentary information to eQTLs in interpreting GWAS hits, especially near functionally important genes or TSS-distal loci.

In total, this work demonstrates the utility of caQTL (and other epigenetic QTL) discovery as a means of uncovering the mechanisms of genetic regulation, alongside eQTL discovery and direct experimental perturbation. In particular, with the increase in availability of molecular data from large, population-scale cohorts, we imagine that a uniform discovery of caQTLs across multiple tissues in the same cohort will be valuable in understanding how consistent the epigenetic regulatory landscape is across tissue types. The relative context-dependence of caQTL discovery compared to eQTL discovery is also worth further investigation – that is, the extent to which context-specific eQTLs are ubiquitously found as caQTLs across developmental stages, circadian rhythm cycles, or stimuli responses. Ultimately, continuing to assay epigenetic molecular phenotypes alongside gene expression will aid in closing the colocalization gap for complex-trait heritability.

## Methods

### Collection and filtering of caQTL summary statistics

caQTL summary statistics were downloaded from ten published studies across different sample groups and tissue types, representing a total sample size of 3,457 (**Supplementary Table 1**). To maintain parity with GWAS which use a threshold of *P*=5×10^−8^, we filtered for SNP-peak pairs with a nominal *P*-value less than 5×10^−8^. We observed that the number of significant single-cell muscle caQTLs from Varshney et al.^29^ far exceeded the counts for other studies; to reduce the bias of our analysis towards muscle caQTLs, we combined the muscle cell “type 1”, “type 2a” and “type 2x” summary stats into a list containing only SNP-peak pairs significant in all three cell types. These three cell types showed very strong sharing of caQTLs in Varshney et al. For SNP selection, we used the list of SNPs passing all filters in Mostafavi et al. (specifically, snp_annotations/filter_snps.txt, **Web Resources**). Briefly, this set started with the list of 13.7 million variants that passed quality control measures for the UKB analyses released by the Neale lab (see “Data availability” in Mostafavi et al.). They further applied the following filters: biallelic autosomal SNP; MAF > 0.01 among the unrelated white British individuals in the UKB; polymorphic in the 1000 Genomes Project phase 3 data. To make the comparisons between GWAS hits and eQTLs consistent, they removed (1) SNPs in LD (r^2^ > 0.8) with predicted protein-truncating or missense mutations, to condition on SNPs putatively acting through gene regulation, and (2) SNPs >1 Mb away from the TSS of any of 18,332 protein-coding genes, which are not tested for eQTLs in GTEx. They further removed SNPs in the major histocompatibility complex region (chr6:28477797-33448354 in hg19). The final filter_snps.txt list contains 6,971,256 SNPs. For full details of SNP filtering, see “Methods: SNP selection” in Mostafavi et al.

### LD clumping to identify lead caQTLs

We replicated the procedure in Mostafavi et al. to clump GWAS/eQTL signals on our caQTL summary stats. For each tissue separately, we used plink’s (**Web Resources**) --clump flag with parameters: *P*-value threshold of 5 × 10^−8^, LD threshold of r^2^ = 0.1 and physical distance threshold of 1 Mb. While Mostafavi et al. used a UKB resource as the LD reference panel, we did not have access to individual level UKB data, so we used 1000G Phase 3 EUR plink files as a reference (**Web Resources**) for all caQTL studies *except* for the DeGorter et al. study, which used LCLs from individuals across six African populations. To clump the LCL caQTLs discovered in African populations, we created an African-only reference panel by downloading the “all_hg38” 2022-08-04 reference set from plink resources, then filtering it for the n=667 individuals with a listed population in [YRI, LWD, GWD, MSL, ESN] (**Web Resources**). After clumping in every tissue, we concatenated the lists of lead caQTLs across tissues to create the “redundant” list of 111,212 lead caQTLs.

We refer to this preliminary list of lead caQTLs as “redundant” because several tissues are represented more than once; for example, bulk liver caQTLs appear in both Wenz et al.^32^ and Currin et al.^22^, neuron caQTLs appear in both Liang et al.^21^ and Zeng et al.^28^, and LCL caQTLs appear in both DeGorter et al.^27^ and Jeong and Bulyk^31^. To reduce the bias in our analysis towards these tissues and cell types, we created a “non-redundant” list of lead caQTLs which included only the highest powered study for each tissue type (see **Supplementary Table 1** for a list of which studies are included). For every duplicated tissue, the highest powered study (with the largest sample size) also had the most lead caQTLs. This non-redundant list included caQTLs from eight of the ten studies, representing a total sample size of 3,204, and consisting of 104,024 lead caQTLs. We used the non-redundant list for all downstream analyses except for tissue-specific analyses, when each tissue in each study in the “redundant” list was analyzed separately.

### GWAS and eQTL lead SNPs

To compare caQTLs to eQTLs and GWAS SNPs, we downloaded the same sets of lead SNPs analyzed by Mostafavi et al. (**Web Resources**, eqtls.all_tissues.clumped.txt and gwas_hits.all_phenos.clumped_sorted.txt) and processed them in the same way. Briefly, this involved applying the same SNP selection filtering as described for the caQTLs, pruning all eQTL brain tissue data except for ‘Brain - Cerebellum’ and ‘Brain - Cortex’, and pruning GWAS traits so that that genetic correlation, *ρ*_g,_ was <0.5 for all trait pairs in the final list (see ‘Methods: Datasets’ in Mostafavi et al. for full details on eQTL/GWAS SNP filtering). This gave us a final list of 118,996 lead eQTLs and 22,119 lead GWAS hits, matching the number reported by Mostafavi et al.

### Collection of SNP and gene features

We downloaded the collection of SNP features (snp_annotations/filter_snps.txt) and gene features (gene_annotations/all_annots_pc_genes.txt) provided by Mostafavi et al. (**Web Resources**). Briefly, these tables contain features for each SNP including distance to TSS, MAF, LD score, and gene density, and features for each gene including length, selective constraint, TSS count, regulatory network connectedness, and GO annotations. See Mostafavi et al. for the full details of how features were collected.

### Calculation of average SNP or gene features

To replicate the analysis of SNPs in Mostafavi et al., we downloaded R scripts provided by the study (**Web Resources**) and adapted them to calculate properties of the lead caQTL list. We also re-ran the code on the eQTL and GWAS lists, getting the same mean results as previously computed. Briefly, this involved computing the mean values of various SNP features in each list of SNPs, or, for gene features, mapping each SNP to the gene with the closest TSS, and then computing the mean feature values of the mapped genes (including duplicated genes – genes mapped to multiple SNPs contributed multiple times to the mean, effectively weighting each gene by the number of independent proximal signals. The full list of gene features analyzed and their computed value for each SNP list can be found in **Supplementary Table 2**. See ‘Methods: Statistical methods’ in Mostafavi et al. for full details, including sources for SNP and gene properties).

### Regression coefficients for gene features

As in Mostafavi et al., to account for correlations between a SNP’s features and the features of its nearest gene, we ran logistic regressions to calculate effect coefficients (e.g. **Fig. 3B**). These regressions estimate the relative effects of gene properties when classifying QTLs/GWAS hits from random SNPs, conditioned on other properties. Specifically, for each SNP list, we selected 100,000 random SNPs from the full set of filtered SNPs, and created an indicator variable to be ‘1’ for SNPs in the list, and ‘0’ for random SNPs. We then ran a logistic regression to predict this indicator variable using normalized gene features of each SNPs nearest gene. As covariates, we included MAF, LD score, gene density, absolute distance to nearest TSS, total gene length, total length of gene coding sequence, as well as dummy variables for 20 quantiles of MAF, LD score, gene density and absolute distance to the nearest TSS. For each SNP type, we first ran this regression using *all* measured gene features as predictor variables in a “conditional” model. The full list of conditional coefficients can be found in **Supplementary Fig. 1** and **Supplementary Table 2.** We then ran a separate “marginal” model for each gene feature to measure the effect of that feature conditioned only on the covariates, rather than on the other gene features. The full list of marginal coefficients can be found in **Supplementary Fig. 2** and **Supplementary Table 2**.

### Creating bootstrapped samples of SNPs to compute confidence intervals

In the same fashion as Mostafavi et al., for each of the three SNP lists, we created 1000 lists of bootstrapped SNPs. Briefly, to create a single bootstrapped list, we used a two-step sampling procedure. First, we sampled tissues (or traits, for GWAS) with replacement, and concatenated the set of SNPs corresponding to each sampled tissue to create an “intermediate list”. We then sampled with replacement from the set of independent LD blocks^52^, and concatenated the set of SNPs from the intermediate list contained in the sampled LD blocks. We then repeated this process 1000 times. For all 1000 lists of bootstrap SNPs, we computed mean values/regression coefficients for SNP and gene features as described above, resulting in 1000 “bootstrap” mean values for each feature. We then report the empirical confidence interval (as the range between 2.5th and 97.5th percentiles) for each feature.

### Creating matched sets of SNPs as a control

In the same fashion as Mostafavi et al., for each of the three SNP lists, we matched every SNP to 1000 control SNPs to create 1000 lists of matched SNPs. Briefly, for every SNP, we extracted SNPs with similar gene density, MAF, and LD score, and sampled from that set 1000 times. For all 1000 lists of matched SNPs, we computed mean SNP and gene features as described above, resulting in 1000 “matched” mean values for each feature. We then report the overall mean of these values and the empirical confidence interval (as the range between 2.5th and 97.5th percentiles) for each feature.

### Calculating *P*-values for mean properties

For all SNP or gene properties, *P*-values were computed on the difference between the mean property for QTLs/GWAS hits and the mean property for control SNPs, under the null hypothesis that the difference is zero. To empirically estimate the null distribution of differences, we took the 1000 bootstrap mean values and 1000 matched mean values described above and calculated the pairwise difference between them, resulting in 1000 null difference values. We then assumed that these null values were normally distributed and divided the observed difference by the standard deviation of the null values to get a z-score. The *P*-value was then computed using a two-sided z-test. Despite using similar scripts and source data provided by Mostafavi et al., our matched confidence intervals were smaller than in the original publication, resulting in more significant *P*-values, although all directions remain unchanged.

### Parker lab QTL analysis

To directly compare caQTLs with eQTLs discovered in the same set of samples, we utilized QTL summary statistics in muscle cells from the Parker lab^29^. The muscle cell types included were Type 1, Type 2a, Type 2x, smooth, satellite, endothelial, T-cell, neuro-muscular junction (NMJ), and fibro-adipogenic progenitors (FAPs). We filtered and clumped the summary statistics for caQTLs and eQTLs in each cell type as described above, then concatenated the lists of QTLs in each cell type to create a “muscle lead caQTL” list (53,958 caQTLs) and a “muscle lead eQTL” list (3,552 eQTLs). We then created matched and bootstrapped sets of SNPs for both of these lists, and calculated gene/SNP properties, confidence internals, and *P*-values as described above. The five cell types listed in **Supplementary Fig. 4A** were the only cell types with more than 100 nominally significant lead caQTLs and eQTLs after filtering.

### Modeling discovery of QTLs and GWAS hits

We leave a full explanation of discovery modeling and visualization of the discovery regions to the **Supplementary Note**. Briefly, our model considers the effect of a single SNP on the accessibility of a single peak (or cRE), which affects the expression of a single gene, in turn affecting a single phenotypic trait. The model assumes no pleiotropy – that is, it assumes all effects of the SNP on the trait are mediated through gene expression, and all effects of the SNP on gene expression are mediated through chromatin accessibility (**Fig. 5A**). To maintain consistency with the model described in Mostafavi et al., we use *β* to denote the effect of the SNP genotype on gene expression, and *γ* to denote the effect of gene expression on the trait. Here, we split the effect *β* into *β*_1,_ the effect of the SNP on chromatin accessibility, and *β*_2,_ the effect of accessibility on gene expression, so that *β* = *β*_1_*β*_2._ We model all effects as linear and all errors as normally distributed:

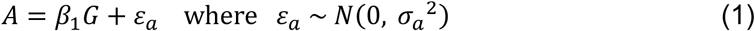

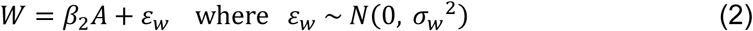

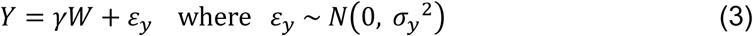

where *G* is the number of non-reference alleles (genotype), *A* is the normalized peak accessibility, *W* is the normalized gene expression, and *Y* is the normalized quantitative trait. The likelihood that a SNP is discovered as a caQTL, eQTL, or GWAS hit (using a fixed significance threshold) then depends on these effect sizes (*β*_1,_ *β*_2,_ *γ*), the study sample size, and the minor allele frequency *p*. In the absence of selection, we model *p* as being uniformly distributed on [0,1], while under selection, we model it using a symmetric Beta distribution negatively correlated with *β*^2^*γ*^2^ (the effect of the SNP on phenotype):

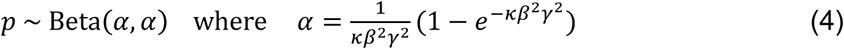

where *κ* is a parameter controlling the strength of selection. For visualization, we set *κ* = 750, although the qualitative shapes of the discovery regions do not depend on the value. We then analytically derive the inequalities under which the single SNP is expected to be discovered in different studies (i.e. as an eQTL, caQTL, or GWAS hit) in terms of *β*^2^ and *γ*^2^ (**Supplementary Note**). The discovery regions in **Fig. 5** use these expressions to show the values of *β*^2^ and *γ*^2^ for which the linear regression in a particular QTL or GWAS discovery is expected to pass the standard significance threshold. Note that because all variances are standardized, the squared effect sizes (*β*_1_^2^, *β* _2_^2^, *γ*^2^) represent *r*^2^ values, the proportion of variance in the outcome variable (*A*, *W*, or *Y*) explained by variance in the causal variable (*G*, *A*, or *W*). Full details can be found in the **Supplementary Note**. See **Web Resources** for links to interactive plots where all model parameters can be adjusted to visualize how they affect the discovery regions.

## Acknowledgments

We thank all members of the Grant and Almasy labs for their feedback on this project. We thank Iain Matheson, Klaus H. Kaestner, and Alexis Battle for sitting on the thesis committee of M.F.D and for providing feedback on this project. M.F.D. is supported by the National Science Foundation Graduate Research Fellowship Program (NSF GRFP). B.F.V. gratefully acknowledges support from the NIH (UM1 DK126194, U24 DK138512, P30 ES013508, R01 DK140340). L.A. is funded by NIAAA U10 AA008401. S.F.A.G. is funded by UM1 DK126194, R01 HD056465 and the Daniel B. Burke Endowed Chair for Diabetes Research.

## Author contributions

M.F.D. conceptualized the project, performed the data analysis, and wrote the manuscript. B.M.W. assisted in curating the collection of caQTLs. B.F.V., L.A., and S.F.A.G. reviewed the manuscript and contributed to the statistical methodology. All authors read and approved the final manuscript.

## Web Resources

- Mostafavi et al. data: https://zenodo.org/records/6618073
- Mostafavi et al. code (commit 0b1002f): https://github.com/hakha-most/gwas_eqtl
- LD clumping:

◦ plink (v1.07) https://zzz.bwh.harvard.edu/plink/download.shtml
◦ 1000G Phase 3 EUR plink reference (1000G_Phase3_plinkfiles.tgz): https://console.cloud.google.com/storage/browser/broad-alkesgroup-public-requester-pays/LDSCORE
◦ 1000G Phase 3 all_hg38 plink reference (used for African reference panel): https://www.cog-genomics.org/plink/2.0/resources#phase3_1kg
◦ 1000G Phase 3 all_hg38 sample data (FTP, contains ancestry info): ftp://ftp.1000genomes.ebi.ac.uk/vol1/ftp/technical/working/20130606_sample_info/20130606_sample_info.xlsx
- Interactive discovery regions from model:

◦ Fig. 5B (neutrality, fixed *β*_2_^2^): https://www.desmos.com/calculator/wkdzzofhmh
◦ Fig. 5C (selection, fixed *β*_2_^2^): https://www.desmos.com/calculator/8vcaemvltw
◦ Fig. 5D (selection, fixed *β*_1_^2^): https://www.desmos.com/calculator/yxwykfsois

## Data and code availability

All code used in this analysis, including the scripts adapted from Mostafavi et al., will be available upon publication.

## Supplementary Figures

**Supplementary Figure 1.**
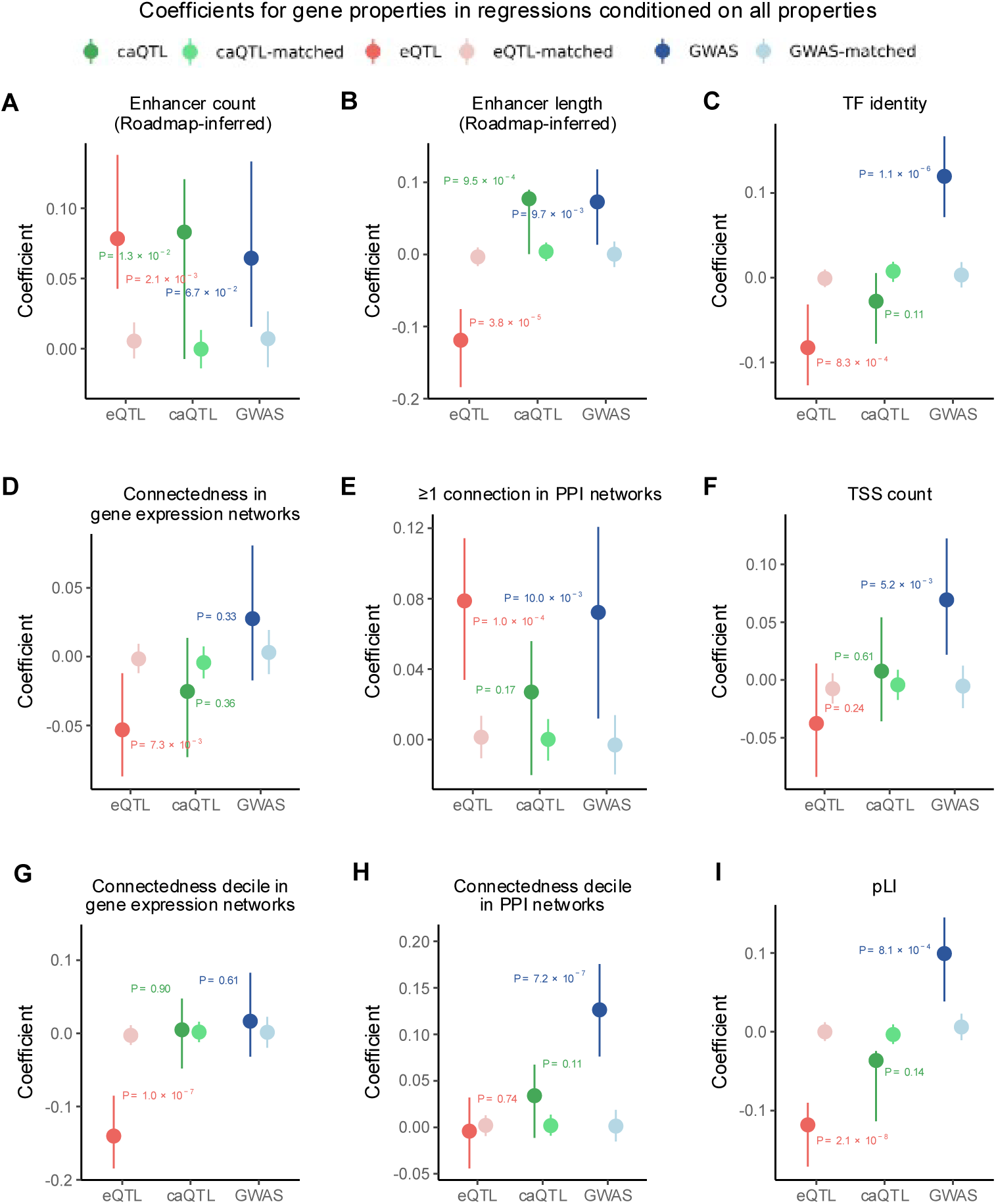
Conditional logistic regression coefficients. Coefficients in a logistic regression framework used to differentiate SNPs in each SNP list from random SNPs, using properties of their nearest gene. For each SNP list, a regression was run which included all gene properties in the same regression, in addition to SNP properties as covariates (**Methods**). Coefficients are shown for (A,B) the count and total length of enhancers inferred through Roadmap data, (C) a binary variable indicating if the gene is a TF, (D) number of neighbors in a co-expression network from Saha et al.^46^, (E) a binary variable indicating the gene has at least one connection in a protein-protein interaction (PPI) network from InWeb^53^, (F) the number of promoters used, (G,H) the decile of connectedness in co-expression and PPI networks, respectively, (I) pLI score. For all panels: confidence intervals for matched lists show 2.5-97.5 percentiles, confidence intervals for lists of discovered SNPs are from bootstrapping. Note that the SNP properties used to match SNPs (TSS distance, LD score, gene density) were included as covariates in the regression, hence coefficients for the matched-SNPs regressions tend to be close to 0. See **Methods** for *P*-value calculations.

**Supplementary Figure 2.**
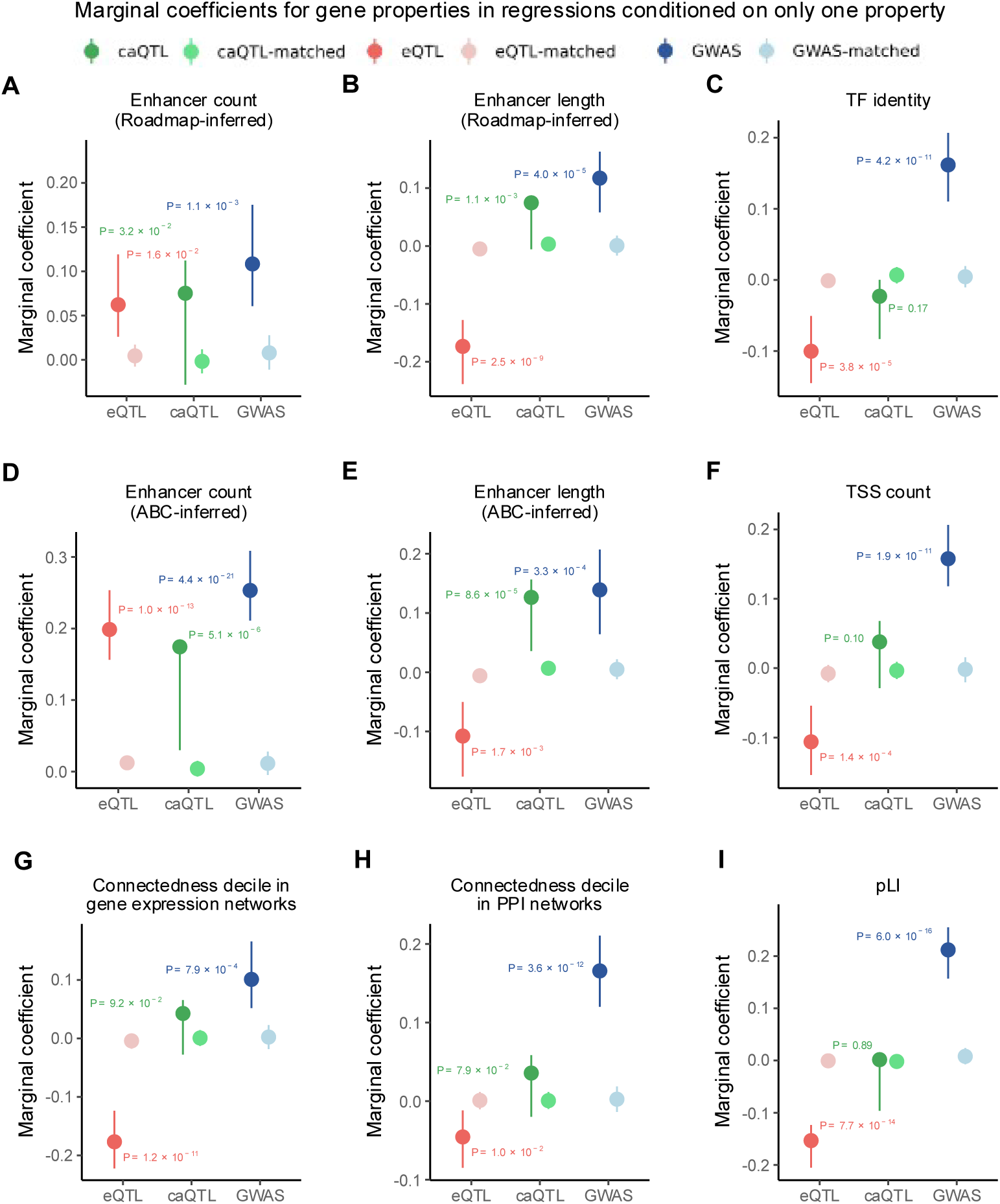
Marginal logistic regression coefficients. Marginal coefficients in a logistic regression framework used to differentiate SNPs in each SNP list from random SNPs, using properties of their nearest gene. For each SNP list and each gene property, a “marginal” regression was run which included only that gene property (unless otherwise specified), in addition to SNP properties as covariates (**Methods**). Coefficients are shown for (A,B) the count and total length of enhancers inferred through Roadmap data (both Roadmap variables were included in the same regression) (panel B is identical to Fig. 3B), (C) a binary variable indicating if the gene is a TF, (D,E) the count and total length of enhancers inferred through the activity-by-contact (ABC)^15^ model (both ABC variables were included in the same regression), (F) the number of promoters used, (G,H) the decile of connectedness in co-expression and PPI networks, respectively, (I) pLI score. For all panels: confidence intervals for matched lists show 2.5-97.5 percentiles, confidence intervals for lists of discovered SNPs are from bootstrapping. Note that the SNP properties used to match SNPs (TSS distance, LD score, gene density) were included as covariates in the regression, hence coefficients for the matched-SNPs regressions tend to be close to 0. See **Methods** for *P*-value calculations.

**Supplementary Figure 3.**
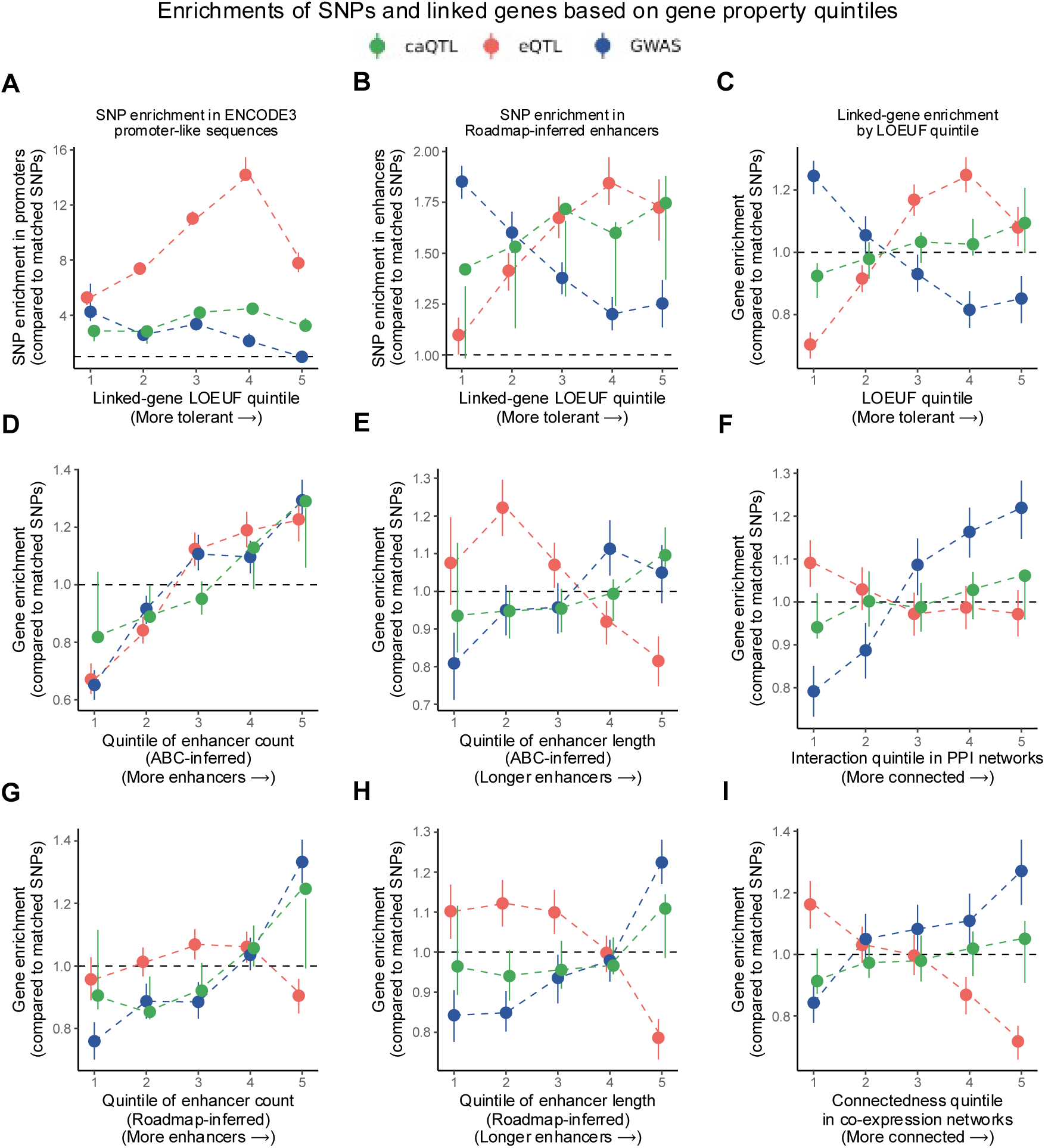
Enrichment of SNPs in gene quintiles. (A,B) GWAS/QTL enrichment compared to matched SNPs, in promoter-like sequences from ENCODE3 (A) and enhancers inferred from Roadmap data (B). For each annotation, the enrichment value is computed as the fraction of SNPs in the annotation divided by the fraction of matched SNPs in the annotation. (C-I) Enrichment of GWAS/QTL genes compared to genes from matched SNPs in gene bins ranked by various gene properties: (C) LOUEF score, (D,E) count and length of ABC-inferred enhancers, (F) connectedness in PPI networks, (G,H) count and length of Roadmap-inferred enhancers, (I) connectedness in co-expression networks (identical to Fig. 3C). Gene enrichment in each bin is calculated as the proportion of GWAS/QTL genes in that bin, divided by the proportion of matched SNP genes in that bin. Confidence intervals are calculated as the bootstrapped upper/lower values divided by the matched SNP 2.5-97.5 percentiles, respectively.

**Supplementary Figure 4.**
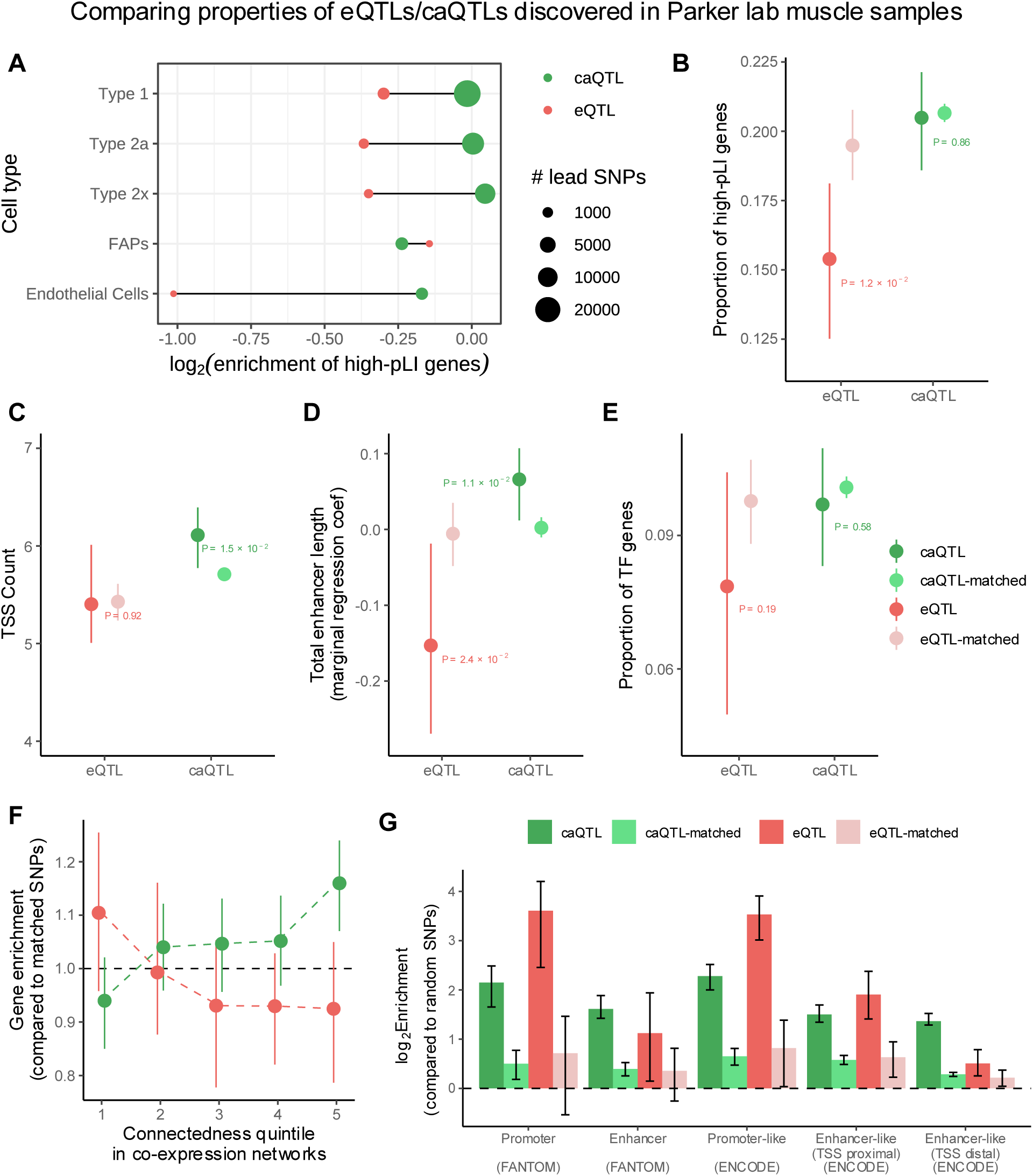
Comparison of caQTLs and eQTLs discovered in the same set of samples. Comparison of eQTLs and caQTLs both discovered in the same set of muscle samples by the Parker lab^29^. (A) Enrichment of high pLI (>0.9) genes in genes mapped by eQTLs and caQTLs discovered in five muscle cell types. (B) Analog of Fig. 2A. (C) Analog of Fig. 3A. (D) Analog of Fig. 3B. (E) Analog of Fig. 3D. (F) Analog of Fig. 3C. (G) Analog of Fig. 4B.

