## Supplementary Note for "A meta-analysis of chromatin-associated loci provides insights into mechanistic interpretations of trait heritability"

The goal of this document is to explain the reasoning behind the original discovery model in Mostafavi et al., and how we expanded it to include a chromatin accessibility component. We try to remain as consistent as possible with the notation and modeling assumptions used in the original model. Similar to Mostafavi et al., the goal of this model is not to predict the discoverability of SNPs under a specific set of parameters, but rather to visualize *qualitatively* how discoverability is related to regulatory effect sizes under natural selection. As such, the model uses simplifying assumptions to make the plotting of discovery regions tractable.

### 1 Discovery model

The model in Mostafavi et al. assumes "that all genetic effects on phenotypes are mediated via cis-effects on gene expression":

$$\text{genotype} \xrightarrow{\beta} \text{gene expression} \xrightarrow{\gamma} \text{phenotype} \tag{1}$$

"where $\beta$ is the per-allele effect size of genotype on expression, and $\gamma$ is the effect size of a unit change in expression on the phenotype." Specifically, they consider linear regression models:

$$W = \beta G + \epsilon_w \tag{2}$$

$$Y = \gamma W + \epsilon_y \tag{3}$$

where $W$ is the expression level of a single gene, $Y$ is a quantitative phenotype, $G$ is the (additive) genotype, $\epsilon_w \sim N(0, \sigma_w^2)$ captures the effect of environment and other causal SNPs (genetic background), and $\epsilon_y \sim N(0, \sigma_y^2)$ captures the effect of environment and other causal genes, which may also be genetically regulated. Combining these models gives:

$$\begin{aligned} Y &= \gamma(\beta G + \epsilon_w) + \epsilon_y \\ &= \beta\gamma G + \epsilon \end{aligned} \tag{4}$$

where $\epsilon = \gamma\epsilon_w + \epsilon_y \sim N(0, \gamma^2\sigma_w^2 + \sigma_y^2)$. This means "the net phenotypic effect of the variant on phenotype is therefore $\beta\gamma$." Note that the dependence of $\text{Var}(\epsilon)$ on $\gamma$ may be problematic, but later they make a simplifying assumption which eliminates this issue.

The genotype $G$ is also modeled as a random variable:

$$G \sim \text{Binomial}(n = 2, p) \tag{5}$$

$$V_p := \text{Var}(G) = 2p(1 - p) \tag{6}$$

where $p$ is the allele frequency, also a random variable. Under neutrality (no selection), the allele frequency is uniform and independent of effect sizes ($p \mid \beta, \gamma \sim \text{Uniform}(0, 1)$), but under selection it is negatively correlated with $\beta^2\gamma^2$. Thus $\text{Var}(G)$, which Mostafavi et al. refers to as "$V_p$", is also random.

#### 2 Discovery thresholds

Consider estimating the eQTL effect $\beta$ using an OLS regression on (2). The estimated effect $\hat{\beta}$ is a random variable which depends on the collected data - the genotype $(g_i)$ and expression level $(w_i)$ for each sample $i$:

$$\hat{\beta} = \frac{\sum_{i=1}^{n}(g_i - \bar{g})(w_i - \bar{w})}{\sum_{i=1}^{n}(g_i - \bar{g})^2} \tag{7}$$

and its variance can be calculated as:

$$\mathsf{Var}(\hat{\beta}) = \frac{\sigma_w^2}{\sum_{i=1}^{n}(g_i - \bar{g})^2} = \frac{\mathsf{Var}(\epsilon_w)}{n\mathsf{Var}(G)} \approx \frac{\mathsf{Var}(W)}{n\mathsf{Var}(G)} \tag{8}$$

Here they make the simplifying assumption $\mathsf{Var}(\epsilon_w) \approx \mathsf{Var}(W)$. They note that the variance in expression can be partitioned into variance explained by the SNP, and variance explained by other factors:

$$\begin{aligned} \mathsf{Var}(W) &= \mathsf{Var}(\beta G + \epsilon_w) \\ &= \beta^2 \mathsf{Var}(G) + \mathsf{Var}(\epsilon_w) \end{aligned} \tag{9}$$

In fact, the proportion of variance in expression explained by the SNP is defined as the expression heritability of the SNP:

$$h^2_{\mathsf{SNP},w} := \frac{\beta^2 \mathsf{Var}(G)}{\mathsf{Var}(W)} \tag{10}$$

The assumption that they make here is that the heritability for a single SNP is small, or equivalently, $\beta^2\mathsf{Var}(G) \ll \mathsf{Var}(\epsilon_w)$. This means that almost all the variance in expression is captured by the noise term: $\mathsf{Var}(W) \approx \mathsf{Var}(\epsilon_w)$.

The estimated effect size is significant if the (squared) z-score is large enough:

$$z = \frac{\hat{\beta}}{\sqrt{\mathsf{Var}(\hat{\beta})}} \quad \longrightarrow \quad z^2 = \chi^2 = \frac{\hat{\beta}^2}{\mathsf{Var}(\hat{\beta})} = \frac{n\mathsf{Var}(G)\hat{\beta}^2}{\mathsf{Var}(W)} > \chi_c^2 \tag{11}$$

Note that the $z^2$ score follows a standard df=1 $\chi^2$ distribution, and so the effect is significant if $\chi^2 > \chi_c^2$, where $\chi_c^2$ is the significance threshold based on $P$-value. For example, the conventional GWAS threshold of $P = 5 \times 10^{-8}$ corresponds to $\chi_c^2 = 29.7$. We expect a SNP to be discovered as an eQTL if $\mathsf{E}[\chi^2] > \chi_c^2$. Since $\mathsf{Var}(W)$ and $n$ are constant for a particular regression, we can define the "study-specific" threshold:

$$c^*_{\mathsf{eQTL}} := \frac{\chi_c^2 \mathsf{Var}(W)}{n_{\mathsf{eQTL}}} \tag{12}$$

Therefore, we expect the variant to be discovered if $\mathsf{E}[\chi^2] > \chi_c^2$, or equivalently:

$$\mathsf{E}\big[\mathsf{Var}(G)\hat{\beta}^2\big] = \mathsf{E}\big[2p(1-p)\big]\beta^2 > c^*_{\mathsf{eQTL}} \tag{13}$$

$$\mathsf{E}\big[h^2_{\mathsf{SNP},w}\big] > \frac{\chi_c^2}{n_{\mathsf{eQTL}}} \tag{14}$$

These equations are very similar for GWAS, regressing on (4):

$$\mathsf{E}\big[2p(1-p)\big]\beta^2\gamma^2 > c^*_{\mathsf{GWAS}} \tag{15}$$

$$\mathsf{E}\big[h^2_{\mathsf{SNP},y}\big] > \frac{\chi^2_c}{n_{\mathsf{GWAS}}} \tag{16}$$

The paper argues that while the trait heritability of SNPs is usually much smaller than the *cis*-expression heritability of SNPs ($h^2_{\mathsf{SNP},y} \approx h^2_{\mathsf{SNP},w} \times 10^{-3}$, see eq. 26), this is roughly balanced by the much larger GWAS sample sizes ($n_{\mathsf{GWAS}} \approx n_{\mathsf{eQTL}} \times 10^3$). Thus, they estimate that both assays generally discover the same proportion (10-20%) of all causal variants.

### 3 Effect of selection

For conciseness, Mostafavi defines $V_p := 2p(1-p) = \mathsf{Var}(G)$, the variance in genotype. Recall that a SNP's discovery depends on its expected contribution to expression or phenotype, which is $\mathsf{E}[V_p \mid \beta,\gamma]\beta^2$ for eQTLs and $\mathsf{E}[V_p \mid \beta,\gamma]\beta^2\gamma^2$ for GWAS hits. These values must be above a threshold:

$$\begin{aligned} \mathsf{E}\big[V_p \mid \beta,\gamma\big]\beta^2 &> c^*_{\mathsf{eQTL}} \\ \mathsf{E}\big[V_p \mid \beta,\gamma\big]\beta^2\gamma^2 &> c^*_{\mathsf{GWAS}} \end{aligned} \tag{17}$$

Under neutrality, the allele frequency is independent of sample size:

$$\mathsf{E}\big[V_p \mid \beta,\gamma,\mathsf{neutrality}\big] \propto 1 \tag{18}$$

while under selection, the allele frequency is negatively correlated with $\beta^2\gamma^2$, so that:

$$\mathsf{E}\big[V_p \mid \beta,\gamma,\mathsf{selection}\big] < \mathsf{E}\big[V_p \mid \beta,\gamma,\mathsf{neutrality}\big] \tag{19}$$

Mostafavi et. al explores several ways to model this correlation, but for visualization purposes settles on:

$$\begin{aligned} \mathsf{E}\big[V_p \mid \beta,\gamma,\mathsf{selection}\big]\beta^2\gamma^2 &\propto \kappa(1-e^{-\beta^2\gamma^2/\kappa}) \\ \mathsf{E}\big[V_p \mid \beta,\gamma,\mathsf{selection}\big] &\propto \frac{\kappa(1-e^{-\beta^2\gamma^2/\kappa})}{\beta^2\gamma^2} \end{aligned} \tag{20}$$

where $\kappa = 2.986$ is a parameter that controls the strength of selection. These variance distributions are described proportionally with a constant omitted, because it is not needed to visualize the discovery regions (see below).

### 4 Visualizing discovery thresholds using simulations

To plot the discovery regions in Fig. 6, Mostafavi et al. sampled 10 million pairs of $(\beta,\gamma)$, assuming that they are independent and normally distributed:

$$\beta,\gamma \sim N(0,1) \tag{21}$$

This allowed them to compute an empirical distribution of variance contribution for each of the four scenarios (eQTL/GWAS, neutrality/selection), conditioned on the randomly drawn $\beta$ and $\gamma$. For example, eQTL discovery under selection depends on the distribution of $\mathsf{E}[V_p \mid \beta,\gamma,\mathsf{selection}]\beta^2$. To set the significance thresholds ($c^*$), they assumed that each study is powered to detect 15% of signals:

$$P\Big(\mathsf{E}\big[V_p \mid \beta,\gamma\big]\beta^2 \quad > \quad c^*_{\mathsf{eQTL}}\Big) = 0.15$$

$$P\Big(\mathsf{E}\big[V_p \mid \beta,\gamma\big]\beta^2\gamma^2 \ > \ c^*_{\mathsf{GWAS}}\Big) = 0.15 \tag{22}$$

That is, for each of the four scenarios, they set the $c^*$ values to be the 85th percentile of the corresponding variance distribution. Recall that $c^* \propto 1/n$, and so setting $c^*$ is equivalent to choosing the sample size for a study with a fixed *P*-value threshold. Intuitively, since SNPs contribute more to expression variance than to trait variance, $c^*_{\mathsf{GWAS}}$ will have to be set lower than $c^*_{\mathsf{eQTL}}$ to capture 15% of SNPs. This is equivalent to assuming that GWAS has a sufficiently higher sample size, although the exact amount depends on the relationship between expression and trait heritability.

Note that because these thresholds were based on percentiles, the significance of a simulated SNP does not depend on the scaling of the $\mathsf{E}[V_p \mid \beta,\gamma]$ distribution. Thus, these distributions are only described proportionally. Fig. 6 of Mostafavi et al. then shows the pairs of $(\beta,\gamma)$ for which the expected variance contribution is greater than $c^*$.

### 5 Standardization

If we assume that the expression and quantitative phenotype data is standardized so that:

$$\mathsf{Var}(W) = \mathsf{Var}(Y) = 1$$

then this lets us simplify the discovery equations:

$$\begin{aligned}
\mathsf{E}\big[h^2_w\big] &= \mathsf{E}\big[V_p \mid \beta,\gamma\big]\beta^2 &&> c^*_{\mathsf{eQTL}} = \frac{\chi^2_c}{n_{\mathsf{eQTL}}} \\
\mathsf{E}\big[h^2_y\big] &= \mathsf{E}\big[V_p \mid \beta,\gamma\big]\beta^2\gamma^2 &&> c^*_{\mathsf{GWAS}} = \frac{\chi^2_c}{n_{\mathsf{GWAS}}}
\end{aligned} \tag{23}$$

We've also abbreviated the subscripts here by letting $h^2_{\mathsf{SNP},w} = h^2_w$, etc.

### 6 Estimation of eQTL and GWAS power

This section is a summary of the explanation in Mostafavi et al. which justifies their claim that "GWAS and eQTL studies are generally low-powered at current sample sizes."

Let $h^2_Y$ refer to the total heritability of the trait, and recall that $h^2_{\mathsf{SNP},y} = h^2_y$ refers to the contribution of a single SNP to trait heritability, i.e. the fraction of trait variance explained by a single SNP. Likewise let $h^2_W$ refer to the total *cis*-heritability of a given gene, that is, the fraction of expression variance explained by *cis*-regulatory variants, and recall that $h^2_{\mathsf{SNP},w} = h^2_w$ refers to the fraction of *cis*-expression heritability explained by a single SNP. Studies estimate that the number of causal SNPs for a complex trait is on the order of 10K-100K, so on average:

$$\frac{h^2_y}{h^2_Y} \approx 10^{-4.5} \tag{24}$$

Likewise, the number of *cis* causal SNPs for a gene's expression is on the order of 10, so:

$$\frac{h^2_w}{h^2_W} \approx 10^{-1} \tag{25}$$

For example: typical values for total heritability are $h_Y^2 \approx 0.2$ and $h_W^2 \approx 0.05$, which, plugging into these equations, gives average SNP heritabilities of $h_y^2 \approx 6 \times 10^{-6}$ and $h_w^2 \approx 5 \times 10^{-3}$, meaning that per-SNP heritability for gene expression is around 3 orders of magnitude greater than for a trait:

$$h_y^2 \approx h_w^2 \times 10^{-3} \tag{26}$$

Further, recall that in expectation, discovered eQTLs satisfy $h_w > \chi_c^2/n_{\text{eQTL}}$ and discovered GWAS hits satisfy $h_y > \chi_c^2/n_{\text{GWAS}}$. For GWAS, the conventional threshold of $P = 5 \times 10^{-8}$ corresponds to $\chi_c^2 = 29.7$, while the typical eQTL nominal threshold of $P \approx 2 \times 10^{-4}$ corresponds to $\chi_c^2 \approx 14$. Plugging these in to the equations above shows that the discovery of a SNP with average heritability requires the sample sizes:

$$n_{\text{eQTL}} > \frac{\chi_c^2}{h_w^2} = \frac{\chi_c^2}{h_W^2\left(\frac{h_w^2}{h_W^2}\right)} \approx \frac{14}{0.05 \cdot 10^{-1}} = 2800 \tag{27}$$

$$n_{\text{GWAS}} > \frac{\chi_c^2}{h_y^2} = \frac{\chi_c^2}{h_Y^2\left(\frac{h_y^2}{h_Y^2}\right)} \approx \frac{29.7}{0.2 \cdot 10^{-4.5}} \approx 4.7 \text{ million} \tag{28}$$

Thus, GWAS and eQTL studies are generally low-powered at current sample sizes.

### 7 Visualizing discovery thresholds using analytical expressions

While Mostafavi et al. used simulations to plot the discovery regions, here we attempt to plot the regions in $(\beta^2, \gamma^2)$ space where we expect variants to be discovered, using analytically-derived inequalities instead of simulations. We will be using the standardization from above where we assume that $\text{Var}(W) = \text{Var}(Y) = 1$.

Recall that the genotype $G$ is modeled as the number of non-reference alleles (5) and $V_p = \text{Var}(G)$ (6):

$$G \sim \text{Binomial}(n = 2, p)$$

$$V_p := \text{Var}(G) = 2p(1-p)$$

#### 7.1 Under neutrality

Under neutrality, the allele frequency $p$ is uniformly distributed, regardless of effect sizes:

$$p \mid \beta, \gamma \sim \text{Uniform}(0, 1) \tag{29}$$

For simplicity of notation, we assume that $\beta$ and $\gamma$ are fixed under the model of neutrality, so we use $\text{E}[V_p]$ to represent $\text{E}[V_p|\beta, \gamma]$.

$$\text{E}\big[V_p\big] = \text{E}\big[\text{Var}(G)\big] = \text{E}\big[2p(1-p)\big] = 2\big(\text{E}[p] - \text{E}[p^2]\big) = 2\left(\frac{1}{2} - \frac{1}{3}\right) = \frac{1}{3} \tag{30}$$

However, it would be more convenient if we standardized $G$ so that $\text{E}[V_p] = 1$. So we can imagine instead of using $G$ as the independent variable, we use $G' = G\sqrt{3}$, so that

$$\mathsf{E}\big[V_p\big] = \mathsf{E}\big[\mathsf{Var}(G')\big] = \mathsf{E}\big[\mathsf{Var}(G\sqrt{3})\big] = \mathsf{E}\big[3\mathsf{Var}(G)\big] = 3 * \mathsf{E}\big[\mathsf{Var}(G)\big] = 1 \tag{31}$$

Returning to (9) with our standardizations, we get:

$$\begin{aligned} 1 &= \beta^2 V_p + \sigma_w^2 \\ \mathsf{E}\big[1 \mid \beta, \gamma\big] &= \mathsf{E}\big[\beta^2 V_p + \sigma_w^2 \mid \beta, \gamma\big] \\ 1 &= \beta^2 + \sigma_w^2 \end{aligned} \tag{32}$$

and likewise:

$$1 = \gamma^2 + \sigma_y^2 \tag{33}$$

Since $\sigma^2 > 0$ in both cases, we know that $0 < \beta^2, \gamma^2 < 1$. So now we can use our discovery equations (23) to get the expected discovery conditions for eQTL and GWAS:

$$\beta^2 \; > \; \frac{\chi_c^2}{n_{\mathsf{eQTL}}} \qquad \text{and} \qquad \beta^2\gamma^2 \; > \; \frac{\chi_c^2}{n_{\mathsf{GWAS}}} \tag{34}$$

Now, we can plug in approximate values for $\chi_c^2$ and $n$ based on current sample sizes and significance thresholds:

$$\beta^2 \; \gtrapprox \; \frac{14}{500} = 0.028 \qquad \text{and} \qquad \beta^2\gamma^2 \; \gtrapprox \; \frac{30}{50,000} = 0.0006 \tag{35}$$

See Fig. 2a for this plot.

#### 7.2 Under selection

To quote Mostafavi et al.:

> "A quantitative treatment of the role of selection is beyond the scope of this paper: ... We make simplistic assumptions to illustrate the qualitative effect of selection on variant discovery in GWAS and eQTL assays."

They then go on to show that their key results displayed in the visualization of discovery regions are robust to modeling assumptions. Similarly, our aim is to use a simple model to visualize the general "flattening" effect that selection has on discovery regions, rather than making precise claims about the role of effect sizes and fitness in genetic architecture.

Here, we will pick a non-uniform distribution for the allele frequency $p$ which simplifies the calculations. Recall that under neutrality, $p$ was modeled as being uniformly distributed. This distribution, $\mathsf{Uniform}(0, 1)$, is just a special case $\mathsf{Beta}(1, 1)$ of the Beta distribution. More generally, we can imagine that $p \sim \mathsf{Beta}(\alpha, \alpha)$, where $0 < \alpha < 1$ is a fixed parameter that controls the likelihood of seeing a rare variant, which can depend on $\beta^2\gamma^2$, the effect of the SNP on fitness. A smaller $\alpha$ is associated with a smaller minor allele frequency, which we can see from plots of the probability density functions (pdfs) (Fig. 1):

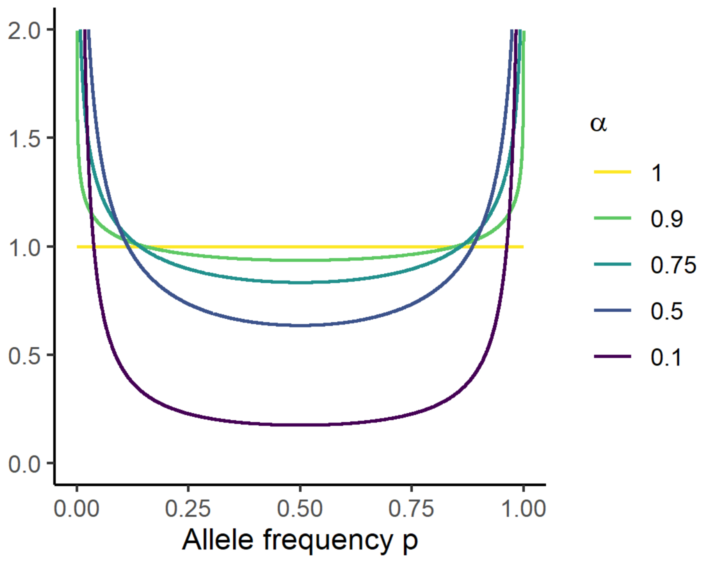


Figure 1: Probability density of Beta$(\alpha, \alpha)$ for different levels of $\alpha$

We can then express the mean and variance of $p$ in terms of $\alpha$:

$$\mathsf{E}[p] = \frac{2\alpha}{\alpha} = \frac{1}{2} \tag{36}$$

$$\mathsf{Var}(p) = \frac{\alpha^2}{(2\alpha)^2(2\alpha+1)} = \frac{1}{4(2\alpha+1)} \tag{37}$$

Then, using the fact that $\mathsf{Var}(p) = \mathsf{E}[p^2] - \mathsf{E}[p]^2$, we can compute the variance in standardized genotype:

$$\begin{aligned}
\mathsf{E}\big[V_p\big] &= 3 * \mathsf{E}\big[\mathsf{Var}(G)\big] \\
&= 3 * \mathsf{E}\big[2p(1-p)\big] \\
&= 6\big(\mathsf{E}[p] - \mathsf{E}[p^2]\big) \\
&= 6\big(\mathsf{E}[p] - \mathsf{Var}(p) - \mathsf{E}[p]^2\big) \\
&= 6\left(\frac{1}{2} - \frac{1}{4(2\alpha+1)} - (\frac{1}{2})^2\right) = \left(\frac{2(2\alpha+1)}{4(2\alpha+1)} - \frac{1}{4(2\alpha+1)} - \frac{2\alpha+1}{4(2\alpha+1)}\right) \\
&= 6\left(\frac{2\alpha}{4(2\alpha+1)}\right) = \frac{3\alpha}{2\alpha+1}
\end{aligned} \tag{38}$$

To model selection, we just need to choose a relationship between $\alpha$ and $\beta^2\gamma^2$ so that $V_p$ is negatively correlated with $\beta^2\gamma^2$. Just like Mostafavi et al., we will introduce a parameter $\kappa$ that controls the strength of selection, with greater values of $\kappa$ corresponding to stronger selection (in Mostafavi et al., greater $\kappa$ corresponded to *weaker* selection, but we feel that the other way is more intuitive, and also simplifies the equations). So, now, letting $F = \kappa\beta^2\gamma^2$ for brevity, we can choose to model the MAF parameter $\alpha$ as:

$$\alpha = \frac{1}{F}(1 - e^{-F}) = \frac{1}{\kappa\beta^2\gamma^2}(1 - e^{-\kappa\beta^2\gamma^2}) \tag{39}$$

This is similar to how Mostafavi et al. chose to model selection (20), but $\mathsf{E}\big[V_p\big]$ looks different:

$$\mathsf{E}\big[V_p\big] = \frac{3\alpha}{2\alpha+1} = \frac{\frac{3}{F}(1-e^{-F})}{\frac{2}{F}(1-e^{-F})+1} = \frac{3(e^F-1)}{e^F(F+2)-2} \tag{40}$$

Note that $F = \kappa\beta^2\gamma^2$ may be thought of as "the effect of the SNP on fitness/selective pressure" if $\kappa$ is thought of as "the effect of the phenotype on fitness". However, we don't interpret $F$ as anything here, and just use it to shorten the equations.

Now we can return to our discovery equations (23) to get the expected discovery conditions for eQTL and GWAS:

$$\frac{3(e^F-1)}{e^F(F+2)-2}\beta^2 \;>\; \frac{\chi_c^2}{n_{\mathsf{eQTL}}} \qquad \text{and} \qquad \frac{3(e^F-1)}{e^F(F+2)-2}\beta^2\gamma^2 \;>\; \frac{\chi_c^2}{n_{\mathsf{GWAS}}} \tag{41}$$

Because the $F$ term depends on $\beta^2\gamma^2$, solving these inequalities is not possible with elementary algebra. But Wolfram Alpha can solve the first (eQTL) inequality if we input it as an equality (letting $c^* = \chi_c^2/n$):

$$\gamma^2 < \frac{c^*_{\mathsf{eQTL}} W\left(\frac{\exp(2-3\beta^2/c^*_{\mathsf{eQTL}})(2c^*_{\mathsf{eQTL}}-3\beta^2)}{c^*_{\mathsf{eQTL}}}\right) - 2c^*_{\mathsf{eQTL}} + 3\beta^2}{c^*_{\mathsf{eQTL}}\kappa\beta^2} \qquad \longrightarrow \qquad \gamma^2 < \frac{W(Be^B)-B}{\kappa\beta^2} \tag{42}$$

where $W(x)$ is the Lambert W function, and we use $B = 2 - 3\beta^2/c^*_{\mathsf{eQTL}}$ to enormously simplify the expression.

To recap, we can plot the following discovery regions, using $x = \beta^2$ and $y = \gamma^2$, and using $B(x)$ as a reminder that $B$ as defined above depends on $\beta^2$:

$$x \;>\; c^*_{\mathsf{eQTL}} = \frac{\chi_c^2}{n_{\mathsf{eQTL}}} \qquad \text{(eQTL, neutrality)} \tag{43}$$

$$xy \;>\; c^*_{\mathsf{GWAS}} = \frac{\chi_c^2}{n_{\mathsf{GWAS}}} \qquad \text{(GWAS, neutrality)} \tag{44}$$

$$y < \frac{W\big(B(x)e^{B(x)}\big) - B(x)}{\kappa x} \qquad \text{where} \quad B(x) = 2 - \frac{3x}{c^*_{\mathsf{eQTL}}} \qquad \text{(eQTL, selection)} \tag{45}$$

$$\frac{3(e^{\kappa xy}-1)}{e^{\kappa xy}(\kappa xy+2)-2}xy > c^*_{\mathsf{GWAS}} \qquad \text{(not closed form)} \qquad \text{(GWAS, selection)} \tag{46}$$

Regardless of whether or not we can find a closed-form solution, we can view the empirical solutions on Desmos.

### 8 Expanding model to include chromatin accessibility

We now consider an expanded model, for the case of one variant, one *cis*-regulatory element, one gene, and one phenotype.

$$\text{genotype} \xrightarrow{\beta_1} \text{chromatin accessibility} \xrightarrow{\beta_2} \text{gene expression} \xrightarrow{\gamma} \text{phenotype} \tag{47}$$

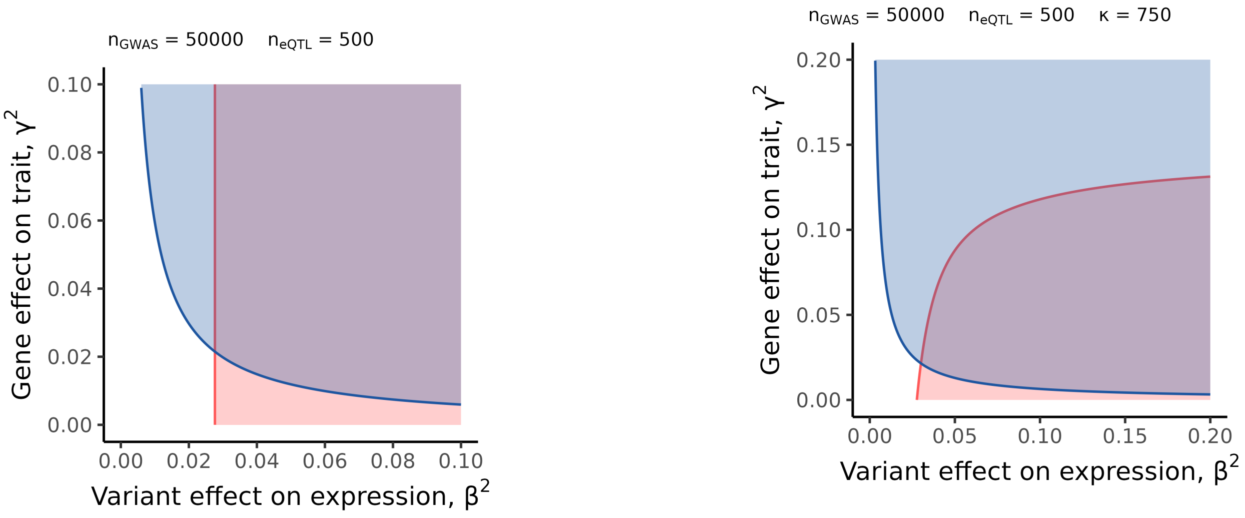


(a) Neutrality

(b) Selection

Figure 2: Discovery regions plotted using expected values (blue=GWAS, red=eQTL)

With regression models:

$$A = \beta_1 G + \epsilon_a \tag{48}$$

$$W = \beta_2 A + \epsilon_w \tag{49}$$

$$Y = \gamma W + \epsilon_y \tag{50}$$

Combining these equations again gives:

$$W = \beta_1 \beta_2 G + \epsilon'_w \tag{51}$$

$$Y = \beta_1 \beta_2 \gamma G + \epsilon'_y \tag{52}$$

where $\epsilon'_w = \epsilon_w + \beta_2 \epsilon_a$ and $\epsilon'_y = \epsilon_y + \gamma \epsilon'_w$. Notice how $\beta = \beta_1 \beta_2$ relates the expanded model to the original model. This new expanded model gives us the discovery requirements for caQTLs:

$$\mathsf{E}\left[h_a^2\right] = \mathsf{E}\left[V_p \mid \beta, \gamma\right] \beta_1^2 \quad > c^*_{\text{caQTL}} = \frac{\chi_c^2}{n_{\text{caQTL}}} \tag{53}$$

#### 8.1 Plotting discovery regions using simulations

Similar to Mostafavi, we sample 10 million values each of $\beta_1, \beta_2, \gamma$, assuming they are i.i.d.:

$$\beta_1, \beta_2, \gamma \sim N(0, 1) \tag{54}$$

We then empirically compute $c^*_{\text{caQTL}}$ as the 85th percentile of the following distribution:

$$\mathsf{E}\left[V_p \mid \beta, \gamma\right] \beta_1^2 \tag{55}$$

where $\mathsf{E}\left[V_p \mid \beta, \gamma\right]$ is calculated under selection using (20).

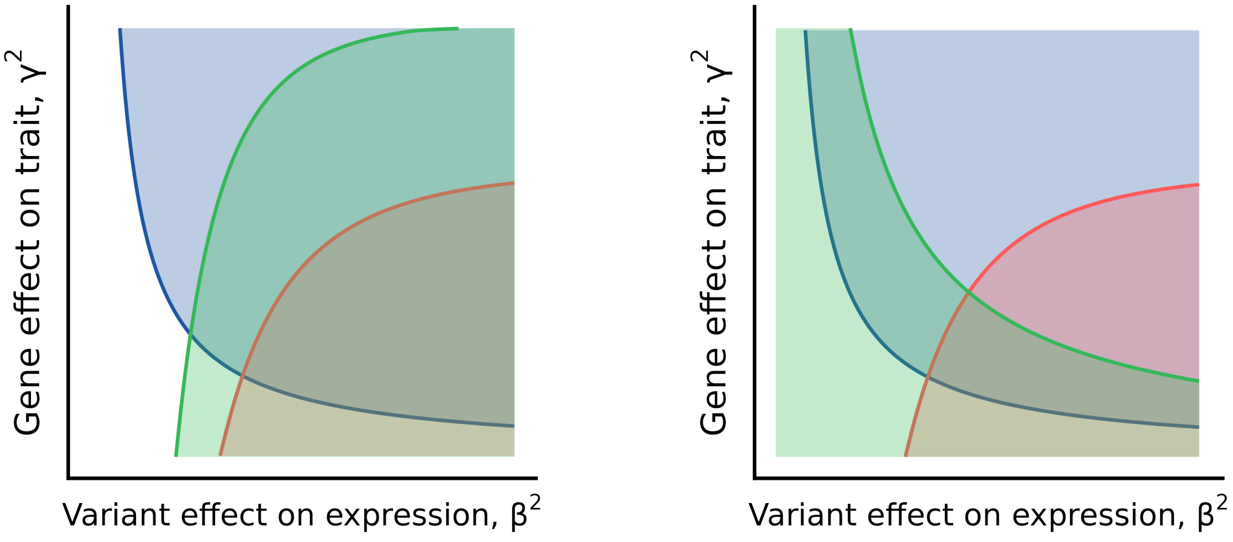


(a) Fixing $\beta_2^2 = 0.45$

(b) Fixing $\beta_1^2 = 2.5$

Figure 3: Discovery regions plotted using simulations (green=caQTL)

Because there are now three parameters, we need to fix either $\beta_1^2$ or $\beta_2^2$ to visualize a slice of the $(\beta^2, \gamma^2)$ space. First, we fix $\beta_2^2$ to the median value of 0.45, producing Fig. 3a. Alternatively, we can fix $\beta_1^2$ to 2.5, producing Fig. 3b. Interestingly, at a power of 15%, $\beta_1^2$ needs to be set to a high value of 2.5 in order for the caQTL region to overlap with the GWAS region, but this could be an artifact of the simplified modeling assumptions, especially the selection equation.

#### 8.2 Plotting discovery regions using expected values

If we fix $\beta_2^2$ to some value, then the caQTL discovery region under neutrality (Fig. 4a) is given by:

$$\beta_1^2 = \frac{\beta^2}{\beta_2^2} > \frac{\chi_c^2}{n_{\text{caQTL}}} = c^*_{\text{caQTL}} \qquad \longrightarrow \qquad \beta^2 > \beta_2^2 c^*_{\text{caQTL}} \tag{56}$$

and under selection, (Fig. 4b):

$$\frac{3(e^F - 1)}{e^F(F+2) - 2} \cdot \frac{\beta^2}{\beta_2^2} > \frac{\chi_c^2}{n_{\text{caQTL}}} = c^*_{\text{caQTL}} \qquad \longrightarrow \qquad \gamma^2 < \frac{W(Ce^C) - C}{\kappa \beta^2} \tag{57}$$

Note that this inequality is almost identical to the eQTL case (41) except we have $\beta_2^2 c^*_{\text{caQTL}}$ in place of $c^*_{\text{eQTL}}$. Thus we can simplify the equation by letting $C = 2 - 3\beta^2/(\beta_2^2 c^*_{\text{caQTL}})$.

Also notice how for the eQTL and caQTL discovery regions to be equal, we must have $n_{\text{caQTL}} = \beta_2^2 \cdot n_{\text{eQTL}}$, meaning for $\beta_2 < 1$ our caQTL sample size can be *smaller* than eQTL sample size while still being expected to capture the same variants (for the set of variants where expression is mediated through accessibility).

On the other hand, we can also fix $\beta_1^2$ to some value, so under selection we have the inequality:

$$\frac{3(e^F-1)}{e^F(F+2)-2}\cdot\beta_1^2 \;>\; \frac{\chi_c^2}{n_{\text{caQTL}}} = c^*_{\text{caQTL}} \qquad\longrightarrow\qquad \frac{3(e^{\kappa}\beta^2\gamma^2-1)}{e^{\kappa}\beta^2\gamma^2(\kappa\beta^2\gamma^2+2)-2} \;>\; \frac{c^*_{\text{caQTL}}}{\beta_1^2} \tag{58}$$

which we once again use WolframAlpha to solve in closed-form in terms of $\gamma^2$, letting $C' = 2 - 3\beta_1^2/c^*_{\text{caQTL}}$ to simplify (Fig. 5):

$$\gamma^2 < \frac{W(C'e^{C'})-C'}{\kappa\beta^2} \tag{59}$$

Note that plotting the caQTL discovery region while fixing $\beta_1^2$ under *neutrality* is pointless, since the discovery under neutrality depends *only* on $\beta_1^2$. This means that the region would either be the whole $(\beta^2, \gamma^2)$ plane, or nothing.

To recap, we can plot the following discovery regions, using $x = \beta^2$ and $y = \gamma^2$, and using $C(x)$ as a reminder that $C$ as defined above depends on $\beta^2$:

$$x \;>\; \beta_2^2 c^*_{\text{caQTL}} \qquad \text{(neutrality, fixed } \beta_2^2\text{)} \tag{60}$$

$$y < \frac{W\big(C(x)e^{C(x)}\big)-C(x)}{\kappa x} \qquad \text{where} \quad C(x) = 2 - \frac{3x}{\beta_2^2 c^*_{\text{caQTL}}} \qquad \text{(selection, fixed } \beta_2^2\text{)} \tag{61}$$

$$y < \frac{W(C'e^{C'})-C'}{\kappa x} \qquad \text{where} \quad C' = 2 - \frac{3\beta_1^2}{c^*_{\text{caQTL}}} \qquad \text{(selection, fixed } \beta_1^2\text{)} \tag{62}$$

Click on the links above to view an interactive graph in Desmos where you can adjust any of the model parameters and see how they affect the discovery regions. However, we stress that the exact values of these parameters should not be over-interpreted as necessary conditions to discover SNPs of particular effect sizes. Rather, the goal of these visualizations is to communicate qualitatively how the relative shape and size of the discovery regions changes as these parameters are adjusted.

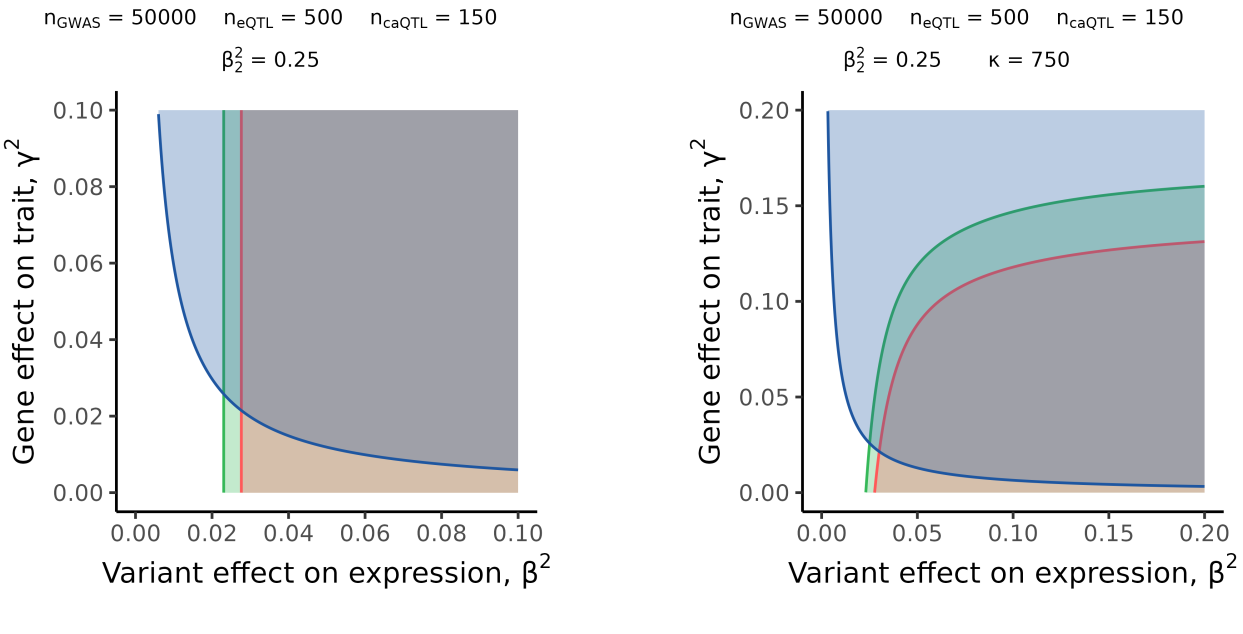


(a) Neutrality

(b) Selection

Figure 4: Discovery regions plotted using equations, fixing $\beta_2^2$ (green=caQTL)

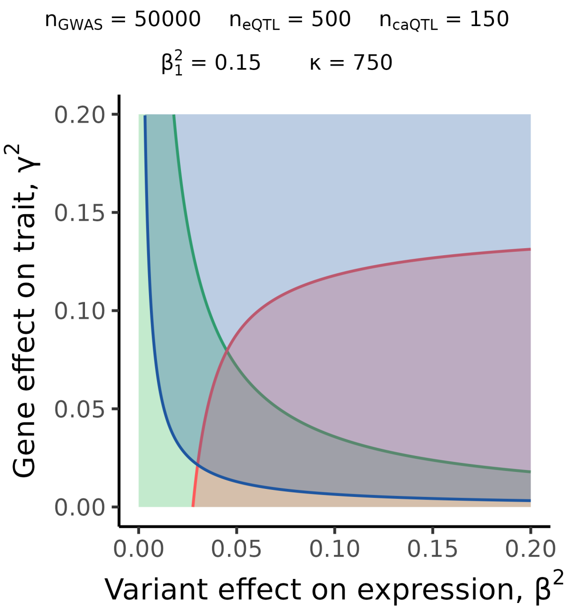


Figure 5: Discovery region under selection, fixing $\beta_1^2$
